# The role of prospective contingency in the control of behavior and dopamine signals during associative learning

**DOI:** 10.1101/2024.02.05.578961

**Authors:** Lechen Qian, Mark Burrell, Jay A. Hennig, Sara Matias, Venkatesh. N. Murthy, Samuel J. Gershman, Naoshige Uchida

**Affiliations:** Department of Molecular and Cellular Biology, Harvard University, Cambridge, MA, USA; Center for Brain Science, Harvard University, Cambridge, MA, USA; Department of Psychology, Harvard University, Cambridge, MA, USA

## Abstract

Associative learning depends on contingency, the degree to which a stimulus predicts an outcome. Despite its importance, the neural mechanisms linking contingency to behavior remain elusive. Here we examined the dopamine activity in the ventral striatum – a signal implicated in associative learning – in a Pavlovian contingency degradation task in mice. We show that both anticipatory licking and dopamine responses to a conditioned stimulus decreased when additional rewards were delivered uncued, but remained unchanged if additional rewards were cued. These results conflict with contingency-based accounts using a traditional definition of contingency or a novel causal learning model (ANCCR), but can be explained by temporal difference (TD) learning models equipped with an appropriate inter-trial-interval (ITI) state representation. Recurrent neural networks trained within a TD framework develop state representations like our best ‘handcrafted’ model. Our findings suggest that the TD error can be a measure that describes both contingency and dopaminergic activity.

## Introduction

The ability to discern predictive relationships between different events is crucial for adaptive behaviors. Early investigations into animal learning revealed that mere contiguity between two events (“pairing”) is insufficient for establishing enduring associations. To understand this, consider Pavlovian conditioning, where an initially neutral cue (conditioned stimulus, CS) is paired with an outcome (unconditioned stimulus, US), such as an electrical shock. Through repeated pairings, animals learn to anticipate the outcome in response to the presentation of just the CS, leading to heightened conditioned responses (e.g., freezing). Now, consider a scenario where the same number of pairings takes place, yet additional shocks occur in the absence of the CS, such that shocks happen with equal likelihood whether or not the CS is present. In such conditions, animals fail to display conditioned responses^1–3^. Moreover, when a CS predicts a decrease in the likelihood of the US, conditioned responses are reduced. Based on these experiments, Rescorla postulated that conditioning depends not on the contiguity between the CS and the US but rather on *contingency* – the degree to which the CS indicates an increase or decrease in the likelihood of the US occurring.

Contingency indicates conditional relationships between different events and is thought to be an important quantity not only in conditioning, but also in causal inference in statistics and artificial intelligence. What is a good measure of contingency, however, remains to be clarified^4–7^. One commonly adopted definition in psychology and causal inference is Δ*P*, the difference in the probability of one event occurring in the presence or absence of another^8–10^. In Pavlovian settings with trial-like structures, such as the present study, Δ*P* can be expressed as Δ*P* = *P*(*US*|*CS*+) − *P*(*US*|*CS*−), where ‘CS+’ and ‘CS-’ signify the presence and absence of a CS, respectively. While mere association does not inherently imply causality, these associations can give rise to perceived causal relationships, and it has been shown that the contingency (Δ*P*) correlates with its strength^6,11,12^. Although ΔP provides a simple definition, its application necessitates a trial-like structure or defined time intervals within which the probabilities of events such as CS and US can be determined^7,13^. Likewise, some behavioral observations cannot be explained by *ΔP*, leading some to argue against the usefulness of contingency in explaining behavior^14^. As a result, efforts have been made to better define contingency^4–7^.

Following Rescorla’s experiments discussed above, further experiments highlighted the crucial role of surprise in the establishment of associations^15^. To account for this, Rescorla and Wagner (1972) postulated that conditioning is driven by the discrepancy between the actual and predicted outcome (prediction error)^16^. Importantly, this contiguity-based model can explain the contingency degradation experiments described above, assuming that the context acts as another CS, which competes with the primary CS^16^. While this “cue-competition” account is attractive, and potentially replaces the classic contingency-based account, the validity of the cue-competition model remains contested^17–20^.

Like *ΔP*, the Rescorla-Wagner model also assumes a trial-based structure, as it does not consider the timing of events either within or outside a trial. To address this limitation, Sutton and Barto developed the temporal difference (TD) learning algorithm, now a fundamental algorithm in reinforcement learning^21,22^, as a prediction error-based model of associative learning^23,24^. TD learning as a model of associative learning in animals finds support in the striking resemblance observed between the activity of midbrain dopamine neurons and the prediction error (TD error) used in TD learning algorithms^25–29^.

Despite the success of TD learning models in accounting for both associative learning and dopamine signals^25,30^, TD models has received various challenges from alternative models. For instance, a recent study^31^ proposed an alternative model for associative learning and dopamine, called an adjusted net contingency for causal relations (ANCCR) model. As the name implies, the ANCCR model posits contingency as a key driver of associative learning and causal inference. Conventional definitions of contingency as well as TD learning models rely on “prospective” predictive relationships between cues and outcomes, i.e. *P*(*US*|*CS*). By contrast in the ANCCR model, learning is driven by “retrospective” relationships, that is the probability of a stimulus (CS) given the outcome (US), or *P*(*CS*|*US*). The authors argued that ANCCR implements causal inference, and that dopamine signals convey a signal for causal learning (the “adjusted net contingency”), not TD errors. Evidence supporting these ideas came from their experiments in examining dopamine signals in mice^31^ and rats^32^ during Pavlovian tasks in which contingency was manipulated. The validity of ANCCR, as well as interpretations of the data presented in these studies, await further examination.

The concept of contingency lies at the heart of learning predictive relationships. Recent work^33,31^ has raised the novel question of whether associations are learned looking forward (prospectively) or looking backward (retrospectively), and how dopamine is involved in these processes^7,31^. Yet how contingency affects dopamine signals and behavior, as well as how dopamine signals relate to causal inference, remains to be determined. To address these questions, we examined behavior and dopamine signals in the ventral striatum (VS) in mice performing Pavlovian conditioning tasks while manipulating stimulus-outcome contingencies. We show that, contrary to previous claims^31,32^, dopamine signals could be comprehensively explained by TD learning models. Furthermore, we found that dopamine signals primarily reflected prospective stimulus-outcome relationships, and strongly violated predictions of the ANCCR model. We then discuss a conceptual framework for how dopamine signals can be related to contingency and causal inference.

## Results

### Contingency degradation attenuates Pavlovian conditioned responding

To study the effects of contingency in a Pavlovian setting, we developed a task for head-fixed mice in which odor cues predicted a stochastic reward (Fig. 1a, b, c). All mice (*n* = 29), after being water restricted, were first trained on one reward-predicting odor (Odor A) that predicted a reward (9 µL water) with 75% probability and one odor (Odor B) that indicated no reward. In this phase (Phase 1), Odor A trials accounted for 40% of trials, Odor B for 20%, with the remaining 40% being blank trials, in which neither odor nor reward was delivered. The timing of task events (Fig. 1b) was chosen such that the trial length was relatively constant, so we could apply the classic Δ*P* definition to our design.

**Figure 1.**
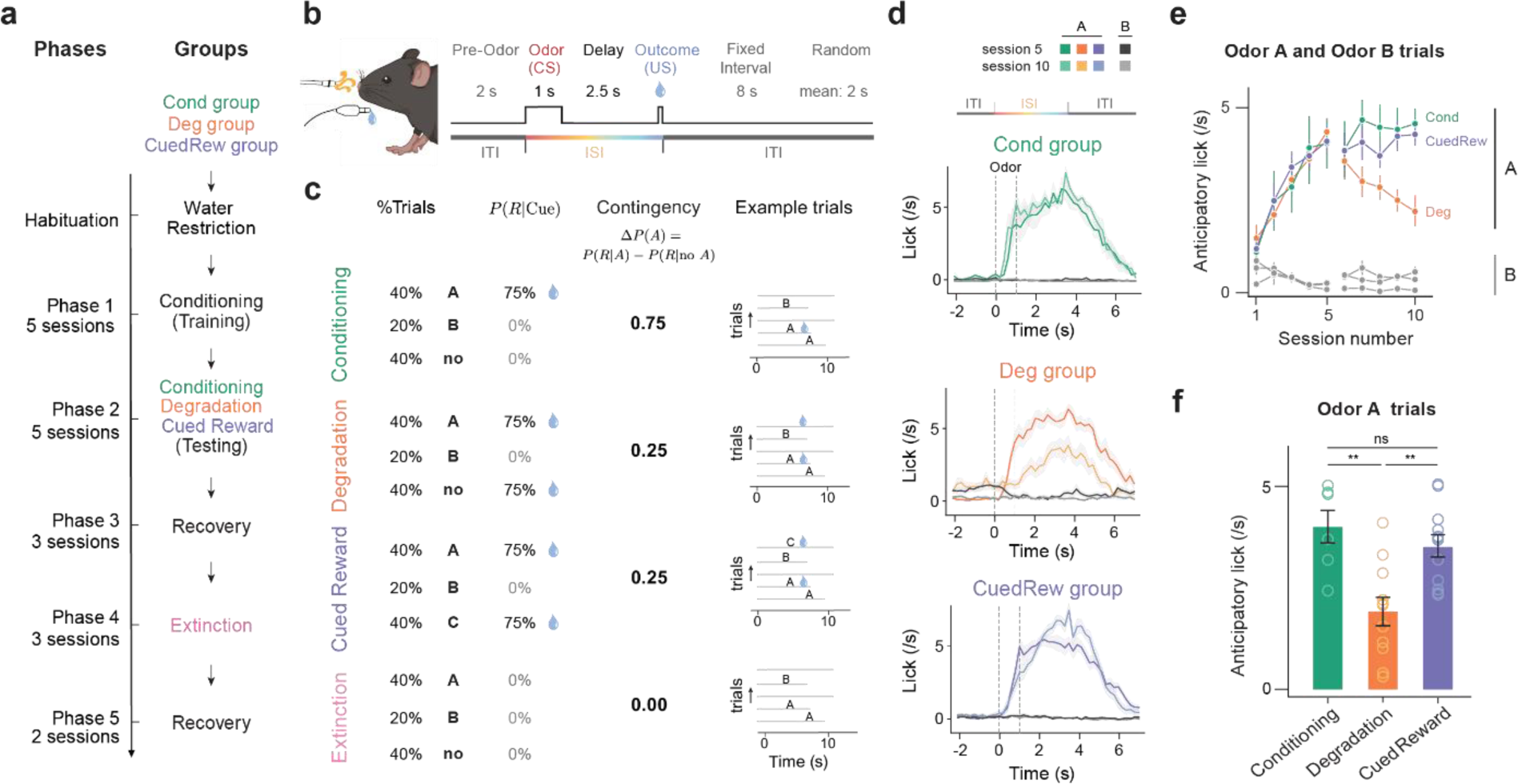
| Dynamic changes in lick response to olfactory cues across different phases of Pavlovian contingency learning task. (a) Experimental design. Three groups of mice subjected to four unique conditions of contingency learning. All animals underwent Phases 1 and 2. Deg group additionally underwent Phases 3-5. (b) Trial timing. (c) Trial parameters per condition. In Conditioning, Degradation and Cued Reward, Odor A predicts 75% chance of reward (9 µL water) delivery, Odor B indicates no reward. In Degradation, blank trials were replaced with uncued rewards (75% reward probability). In Cued Reward, these additional rewards were cued by Odor C. In Extinction, no rewards were delivered. (d) PSTH of average licking response of mice in three groups to the onset of Odor A and Odor B from the last session of Phase 1 (session 5) and Phase 2 (session 10). Shaded area is standard error of the mean (SEM). Notably, the decreased licking response during ISI and increased during ITI in Deg group. (green, Cond group, *n* = 6; orange, Deg group, *n* = 11; purple, CuedRew group, *n* = 12 mice). (e) Average lick rate in 3s post-cue (Odor A or B) by session. Error bars represent SEM. (f) Average lick rate in 3s post Odor A in final session of each condition. Asterisks denote statistical significance: ns, *P* > 0.05; **, *P* < 0.01, Student’s *t*-test, indicating a significant change in licking behavior to Odor A in Deg group across sessions.

In Phase 1, Odor A has positive stimulus outcome contingency, being predictive of reward (R; Fig. 1c). This can be quantified using the commonly applied Δ*P* definition of contingency: e.g., Δ*P*(*A*) = *P*(*R*|*A*+) − *P*(*R*|*A*−) = 0.75 − 0 = 0.75 in Phase 1. Conversely, Odor B has a negative stimulus-outcome contingency: Δ*P*(*B*) = *P*(*R*|*B*+) − *P*(*R*|*B*−) = 0 − 0.375 = −0.375. Consistent with these contingencies, all animals developed anticipatory licking to Odor A, but not Odor B, within five training sessions (Fig. 1e).

In Phase 2, animals were split into groups (Fig. 1a). The first group (‘Cond’, *n* = 6) continued being trained on the identical conditioning task from Phase 1. With no change in contingency, the behavior did not significantly change in a further five sessions of training (Fig. 1d, e).

The second group (‘Deg’, *n* = 11) experienced contingency degradation. To reduce the contingency of Odor A, either *P*(*R*|*A*+) can be decreased or *P*(*R*|*A*−) can be increased. We increased *P*(*R*|*A*−) by introducing uncued rewards, an experimental design termed ‘contingency degradation’^34^. Blank trials from Phase 1 were replaced with ‘background water’ trials in which a reward was delivered on 75% of these trials. In this condition, *P*(*R*|*A*+) remains unchanged at 0.75, while *P*(*R*|*A*−) increases to 0.5 (2 out of every 3 non-Odor A trials are background water trials, of which 75% are rewarded thus *P*(*R*|*A*−) = 2/3 × 0.75 = 0.5). As a result, Δ*P*(*A*) is reduced to 0.25. Concomitant with this decreased contingency, the anticipatory licking to Odor A decreased across five sessions of Phase 2 in the Deg group (*t*_11_ = −4.78, *P* = 0.00074, paired *t*-test). Moreover, Deg group animals increased licking during the inter-trial intervals (ITIs, *t*_11_ = 3.34, *P* = 0.0074, paired *t*-test), potentially reflecting an increased baseline reward expectation. Additionally, the Deg group exhibited longer latencies to initiate licking and an increase in trials where mice did not lick before water delivery in Odor A trials (Extended Data Fig. 1d, e).

The decrease in anticipatory licking, rather than reflecting the decreased contingency, could reflect satiety effects as animals in the Deg group receive twice as many rewards per session as the Cond group. We do not believe satiety explains this effect for at least two reasons: (1) all animals still received and drank about 1 ml supplementary water after each session, and (2) in all but the first degradation session, anticipatory licking was diminished compared to Cond controls in early trials (Extended Data Fig. 1f).

Nevertheless, a third group (‘CuedRew’) was included as a control for satiety effects. This group received identical rewards to the Deg group, but rather than delivering uncued rewards during the previously blank trials, these rewards were delivered following a third odor (Odor C). Unlike animals in the Deg group, animals in the CuedRew group did not decrease anticipatory licking to Odor A. Furthermore, anticipatory licking, background licking and licking latency were similar to the Cond group (Fig. 1d, e; Extended Data Fig. 1).

Δ*P*(*A*) is 0.25 in the Cued Reward condition, for identical reasoning as the Deg group. This indicates that the Δ*P* definition of contingency cannot be the sole determinant of conditioned responding (Fig. 1c). This phenomenon has been previously noted in the behavioral responses in conditioning tasks during contingency degradation^14,35^. It is not resolved by considering a retrospective definition of contingency. Consider Δ*P_retro_*(*A*) = *P*(*A* + |*R*) − *P*(*A* − |*R*) in both the CuedRew and Deg groups, this quantity is identical, with Odor A preceding the reward 50% of the time in both conditions.

In the subsequent stage of our investigation (Phase 3; ‘Recovery 1’), we reinstated the original conditioning parameters for the Deg group, which increased the contingency back to 0.75 for Odor A, yielding an immediate recovery of the level of anticipatory licking (Extended Data Fig. 1g).

To compare the behavior and neural correlates of contingency, we also introduced an Extinction phase (Phase 4) to the Deg group. In this phase, no reward was ever delivered following either odor cue. Over three sessions, anticipatory licking to Odor A gradually waned. Finally, during a second recovery phase (Phase 5; Recovery 2), the anticipatory response to Odor A was effectively reinstated (Extended Data Fig. 1g).

Notably, apart from the Extinction phase, the probability of a reward following Odor A was constant at *P*(*R*|*A*) = 0.75 throughout the experiment while behavior changes considerably. Clearly, the contrast against the probability of reward in the absence of a cue is an important consideration for anticipatory behaviors, with marked changes during contingency degradation. However, the Cued Reward control showed it is not as straightforward as the contrast between the absence and presence of a cue.

### Contingency degradation attenuates dopaminergic cue responses

Given the well-documented role of dopamine in associative learning, we sought to characterize the activity of dopamine neurons in our Pavlovian contingency manipulation task. We monitored axonal calcium signals of dopamine neurons using a multi-fiber fluorometry system^36^ with optical fibers targeting 6 locations within the ventral striatum (VS), including the nucleus accumbens (NAc, medial and lateral) and the olfactory tubercle (OT, 4 locations; Fig. 2a, b). Recordings were made only in the Deg and CuedRew groups, with the final session of Phase 1 used as the within-animal conditioning control. To ensure similar levels of calcium sensor expression across the six recording locations, we employed a transgenic approach by crossing a transgenic mouse line expressing the Cre recombinase in dopamine neurons (DAT-Cre)^37^ and a reporter line that expresses a calcium sensor GCaMP6f in a Cre-dependent manner (Ai148)^38^. Fiber locations were verified using post-mortem histology (see Methods for exclusion criteria, Fig. 2b). All results presented in the main text are from the lateral nucleus accumbens (lNAc), where TD error-like dopamine signals have been observed most consistently^39^, though the main findings are consistent across all locations (minimum cosine similarity between any other area and lNAc’s DA signals during odor A rewarded trials: 0.92, Extended Data Fig. 2).

**Figure 2.**
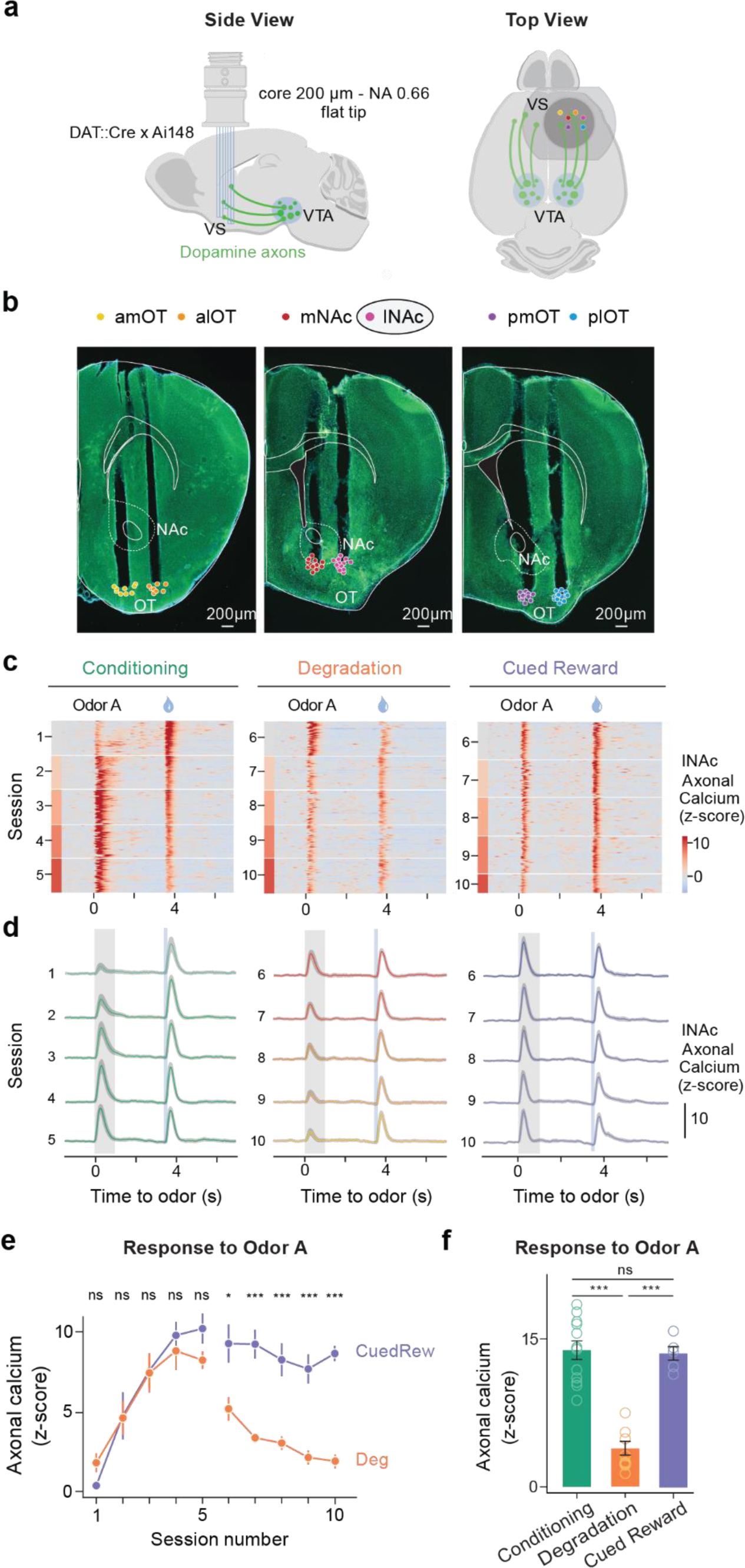
| Dopamine axonal activity recordings show different responses to rewarding cues in Degradation and Cued Reward conditions. (a) Configuration of multifiber photometry recordings. Coronal section from one DAT::cre x Ai148 mouse showing tracts for multiple fibers in the VS. Data recorded from lNAc is used in the following analysis. lNAc, Lateral nucleus accumbens; mNAc, Medial NAc; a lot, anterior lateral olfactory tubercle; plot, posterior lateral OT; amOT, anterior medial OT; pmOT, posterior medial OT. (b) Heatmap from two example mice (mouse 1, left two panels, mouse 2, right panel) illustrating the z-scored dopamine axonal signals in Odor A rewarded trials (rows), aligned to the onset of Odor A for three conditions. (c) Population average z-scored dopamine axonal signals in response to Odor A and water delivery. Shaded areas represent SEM. (d) Mean peak dopamine axonal signal (z-scored) of Odor A response by sessions for the Deg group (orange) and the CuedRew group (purple). Error bars are SEM. *, *P* < 0.05; ***, *P* < 0.001, Welch’s *t*-test. (e) Mean peak dopamine axonal signal (z-scored) for the last session in Phase 1 (Conditioning) and 2 (Degradation and Cued Reward) for both Deg and CuedRew groups. Error bars represent SEM. ns, *P* >0.05; ***, *P* < 0.001, Welch’s *t*-test.

During Phase 1 (initial conditioning) dopamine axons in lNAc initially responded strongly to water and weakly to Odor A (Fig. 2c, d). As learning progressed, the response to water gradually decreased, while the response to Odor A increased over the course of 5 sessions (*t*_13_ = 4.81, *P* = 0.0004, paired *t*-test, cue response first vs. last session of Phase 1), broadly consistent with previous reports of odor-conditioning on stochastic rewards^29,40^.

During contingency degradation (Deg group, Phase 2), the response to Odor A decreased across sessions (*t*_8_ = −11.50, *P* = 8.4 ×10^-6^, paired *t*-test, cue response, session 6 versus 10) consistent with the changes in anticipatory licking and other recent reports of dopamine during contingency degradation^10,31,32^ (Fig. 2e, f). However, in the Cued Reward condition (CuedRew group, Phase 2), the response to Odor A did not decrease compared to the Phase 1 response (*t*_5_ = −1.12, *P* = 0.32, paired *t*-test, cue response first vs. last session of Phase 2), aligning with the behavioral results but conflicting with the idea that dopamine neurons encode contingency, at least so far as defined by Δ*P*.

In the additional phases (3-5) in the Deg group, dopamine also mirrored behavior: the response to Odor A quickly recovered in Recovery 1 (Phase 3), decreased during Extinction (Phase 4) and recovered again during Recovery 2 (Phase 5; Extended Data Fig. 3a). These results show that dopamine cue responses track the stimulus-outcome contingency in our Pavlovian contingency degradation and extinction paradigms although they deviated from the contingency in the CuedRew group. Still, in all groups and phases, dopamine tracked anticipatory licking.

### TD learning models can explain dopamine responses in contingency degradation

In both behavior and dopamine, the responses are not fully explained by contingency: there were diminished responses during contingency degradation, but not when the additional rewards are cued. Given the match between dopamine responses and behavior, rather than consider new definitions of contingency, we sought to test if temporal difference (TD) models, which so far have been highly successful in accounting for dopamine activity, are able to explain the discrepancies from the contingency account.

Dopamine neurons are thought to convey TD errors, denoted by *δ* and defined by the equation: *δ*_*t*_ = *r*_*t*_ + *γV*(*s*_*t*+1_) − *V*(*s*_*t*_), with *r*_*t*_ representing reward at time *t*, *s*_*t*_ representing the state at time *t*, *V*(*s*_*t*_) is the value at state *s*_*t*_, and *γ* is the temporal discount factor (0 < *γ* < 1). Value *V*(*s*_*t*_) is defined as the expected sum of all future rewards starting from time *t*, with each future reward discounted by the factor *γ* at each time step. The role of the TD error in learning is to iteratively refine the value estimate (Fig. 3a), ultimately guiding behavior.

**Figure 3.**
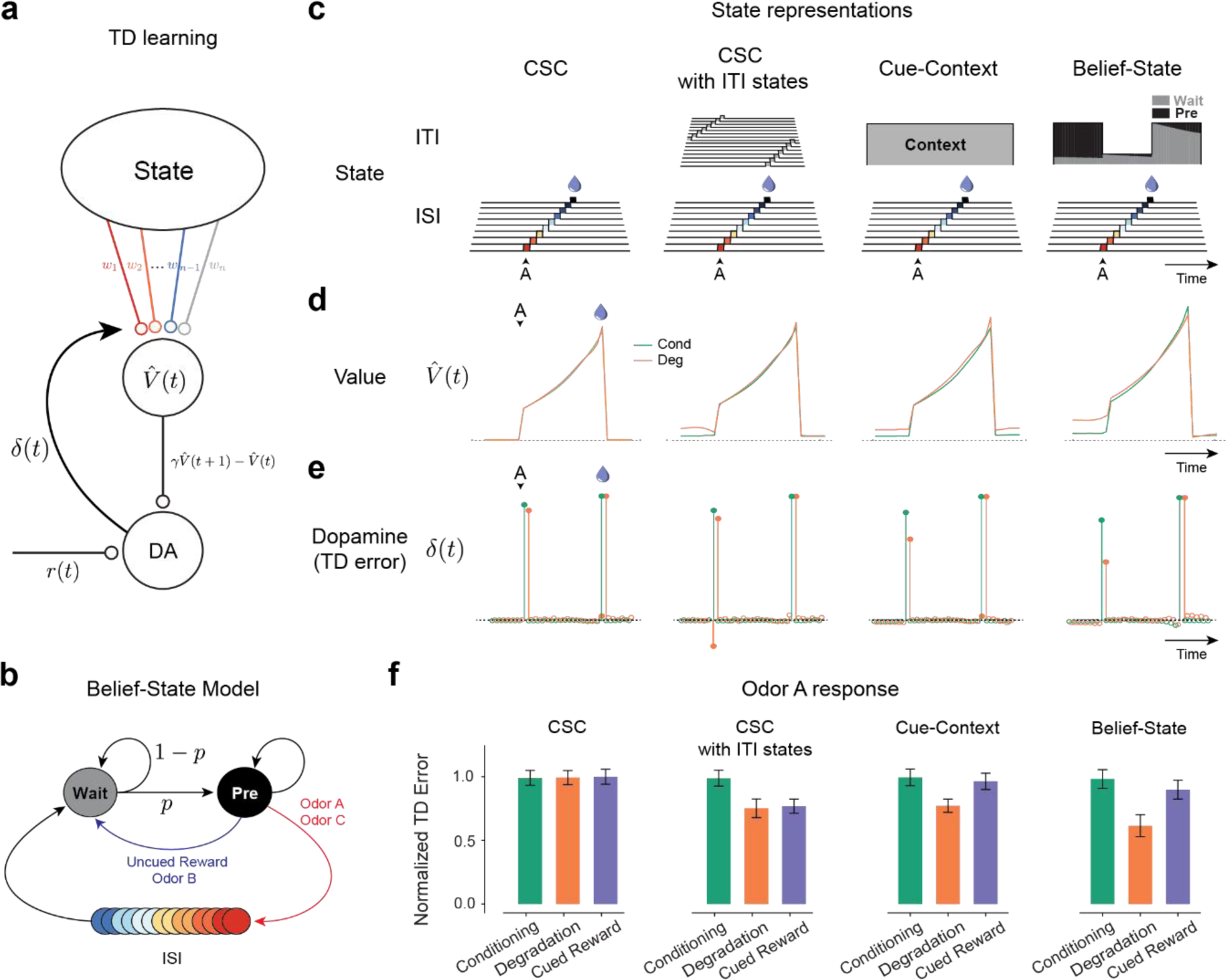
| TD learning models can explain dopamine responses in contingency degradation with appropriate ITI representation. (a) Temporal Difference Zero, TD(0), model – The state representation determines value. The difference in value between the current and gamma-discounted future state plus the reward determines the reward prediction error or dopamine. This error drives updates in the weights. (b) Belief-State Model: After the ISI, the animal is in the Wait state, transitioning to the pre-transition (‘Pre’) state with fixed probability p. Animal only leaves Pre state following the observation of odor or reward. (c) State representations: from the left, Complete Serial Compound (CSC) with no ITI representation, CSC with ITI states, Cue-Context model and the Belief-State model. (d) Value in Odor A trials of each state representation using TD(0) for Conditioning and Degradation conditions (e) TD error is the difference in value plus the reward. (f) Mean normalized TD error of Odor A response from 25 simulated experiments. Error bars are SD.

The response to Odor A differed most between our three test conditions (Conditioning, Degradation, Cued Reward) and thus our modeling efforts initially focused on explaining these changes. Noting there was no reward at the time of Odor A, by the definition of TD error, the response to Odor A is *γV*(*s*_*t*+1_) − *V*(*s*_*t*_), the difference between the value in the state immediately after Odor A (ISI) and the value in the state immediately before Odor A (pre-Odor ITI).

Previous studies have indicated that the ability of TD learning models to explain dopamine responses and conditioned behaviors depends critically on what types of state representations the models use^41–44^. We therefore tested TD learning models (Fig. 3a) equipped with four different types of state representation (Fig. 3c).

In the original application of TD models to dopamine activity, only the interval between a CS and a US, i.e. inter-stimulus interval (ISI), was considered, and was represented using a ‘complete serial compound’ (CSC) representation, sometimes known as a tapped delay line^25,45^. In this construction, a presentation of a stimulus triggers a sequential activation of sub-states, each of which represents a time step after the stimulus (Fig. 3c). At any given time after the stimulus, only one sub-state is active. The value estimate ^*V*^^(*s*_*t*_) is then computed as the weighted sum of these substates which in CSC reduces to be the weight of the active substate.

While this ISI-only CSC state representation is successful in explaining many properties of the dopamine response to conditioned stimuli, it fails to predict the result of our experiments. As the ISI period is identical between conditions and there is no representation of the ITI period, the TD error for Odor A is unchanged between conditions (Fig. 3f).

An extension of this ISI-only model (CSC with ITI states model) models both the ISI and ITI using CSC, resetting with each odor. While this model predicts a decrease for Degradation, it also predicts a decrease in the Cued Reward condition (Fig. 3f), conflicting with our results.

Rather than representing the ITI with many consecutive states, it is possible to represent it as a single state. This model, which we term the Cue-Context model, is functionally similar to the previously developed cue competition model^16–19^. Our Cue-Context model extends the original CSC model with a state that is constantly on (the ‘context’) during both the ISI and ITI (Fig. 3b). This model successfully predicts the pattern of experimental results we observed, with a decrease in the Odor A response during Degradation and a smaller decrease during Cued Reward (Fig. 3f). This can be understood as the context acquiring value, lesser in the Cued Reward condition because the increased value (more rewards) is attributed to both the context and the new odor. On the other hand, in the Degradation condition, the increased value is attributed fully to the context. By increasing the context and thus value during the pre-Odor ITI period, the TD error at Odor A is diminished. While this produces a qualitatively correct pattern of results, it requires a temporal discount factor that is well below previously reported values^46–49^ to produce the quantitatively correct pattern (Extended Data Fig. 4).

We therefore considered whether further information about the experimental design could be used to refine the state representation. Inspired by previous work showing that dopamine neurons are sensitive to hidden state inference in a task with stochastically timed rewards^50,51^, we considered a ‘belief state’ representation, a vector of probabilities for each possible hidden state (Belief-State TD model; Fig. 3b): the ‘Wait’ state, which reflects early ITI (a minimum fixed amount of waiting period in which there is no chance of an uncued reward or odor being delivered), and the ‘pre-transition’ (Pre) state, in which there is an imminent chance of reward or odor being delivered. The transition and observation matrix, which are used to compute the probability of each state, were derived from the experimental settings, assuming a fixed probability of transition from the Wait to Pre state, modeling a growing anticipation of the next trial beginning. Using this state representation improved the quantitative accuracy of the model for a given *γ* versus the Cue-Context, and accurately predicted the experimental data at a value of consistent with previously reported results^46–49^ (Fig 3f., Extended Data Fig. 4).

To test which of these two models, Cue-Context or Belief-State, best describes the state representation driving dopamine responses and behavior, we focused our analysis on the ITI period: whereas the Cue-Context representation models the ITI as a single, homogenous state (the context), the Belief-State model captures temporal heterogeneity by modeling it as the gradual transition between two states – capturing a growing anticipation of the next trial or reward (Fig. 4a).

**Figure 4.**
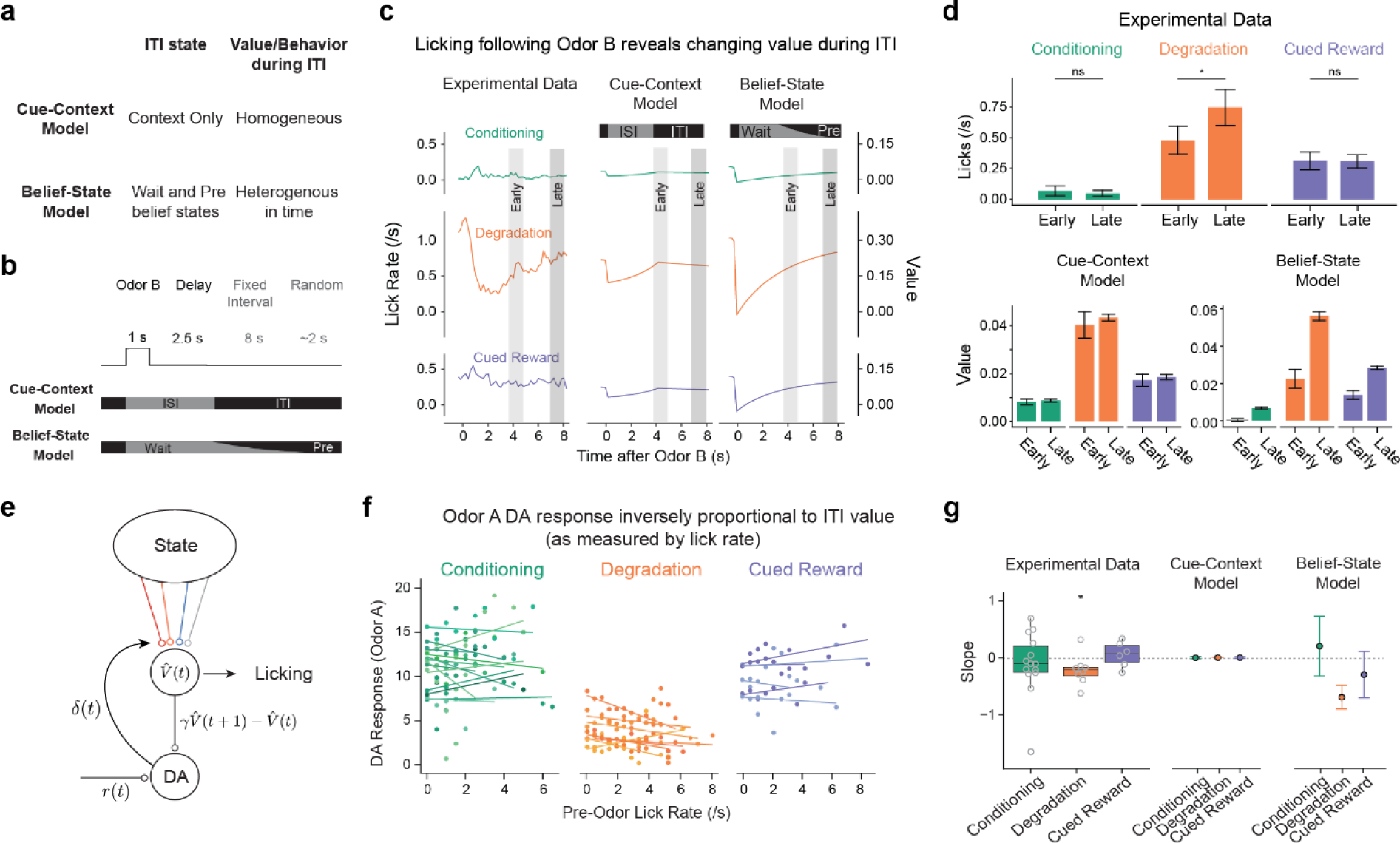
| Belief-State model, but not Cue-Context model, explains variance in behavior and dopamine responses. (a) Cue-Context model and Belief-State model differ in their representation of the ITI. (b) Odor B predicts no reward and at least 10 s before the start of the next trial. (c) Odor B induces a reduction in licking, particularly in the Degradation condition, which matches the pattern of value in the Belief-State model better than the Cue-Context model. (d) Quantified licks (top) from experimental data in early (3.5-5s) and late (7-8s) post cue period. Error bars are SEM, *, *P* < 0.05, paired *t*-test. Value from Cue-Context and Belief-State model for the same time period, error bars are SD. (e) If licking is taken as a readout of value, then ITI licking should be inversely correlated with dopamine. (f) Per animal linear regression of Odor A dopamine response (z-score axonal calcium) on lick rate in 2s before cue delivery. (g) Summarized slope coefficients from experimental data (left) and models (right). Boxplot shows median and IQR, one sample *t*-test.

In Pavlovian settings, anticipatory licking (as opposed to consummatory licking) has been used as a measure of current value – for example, animals will lick more to cues that predict greater rewards^26^. Odor B provides an opportunity to examine whether the ITI is a heterogenous interval. This is because Odor B predicts no reward within the current trial (and thus no consummatory licking) but also provides further information that no odor or uncued reward will be delivered for the length of one trial. In this way, Odor B provides 100% certainty to the animal that while they are in the ‘Wait’ state. Consistent with this understanding of the task structure, the delivery of Odor B during the Degradation condition prompted animals to stop licking. Both the Cue-Context and Belief-State models capture this effect. The crucial difference is how the lick rate recovers. In the Cue-Context model, ITI value is related to a single state, which without reward decreases at the rate of *α* (learning rate). In the Belief-State model, value continually increases (Fig. 4c) across the entire ITI, as the increased belief that the next trial is imminent increases continuously. We find that the lick pattern following Odor B matches the pattern of the Belief-State model and not the Cue-Context model, with a sudden decrease in licking followed by a gradual increase in the Degradation condition that is unrelated to the ISI length (Fig. 4c, summarized in Fig. 4d).

This pattern of licking behavior also suggests that the animals do not develop more complex models of timing. Odor B predicts approximately ten seconds with no reward. An ideal agent would not lick during this time, waiting until the transition to an uncued reward is possible. The mice instead resume licking within 2-3 seconds of Odor B delivery, with the lick rate increasing over several seconds.

While the value from the Belief-State model explains the time course of licking following Odor B, this account does not, by itself, explain the decrease in anticipatory licking in response to Odor A (Fig. 1d). This decreased responding is a consistent feature of contingency degradation^10,34^. We show, in Extended Data Fig 5, that if licks carry a small effort cost and licks are distributed according to relative value, then the Belief-State model can account for the increased licking in the pre-odor period and decreased licking during the ISI.

Having shown that the lick rate is explained by the changing value in the Belief-State model, we wished to test whether this could be used to explain trial-by-trial variance in the dopamine response. Continuing with the assumption that licking is a moment-by-moment measure of value, our Belief-State model predicts there should be an inverse correlation between the pre-odor lick rate and the Odor A dopamine response. To test this, we correlated the number of licks in the two seconds before the cue to the Odor A response on a trial-by-trial basis, regressing a linear model independently for each mouse (pooling the last two sessions of each condition under the assumption that the task was well-learnt in these sessions).

Only in the Degradation condition was there a significant negative correlation between the pre-odor lick-rate and the Odor A dopamine response for the population (Fig. 4f, g). This can be explained by the Belief-State model, as ITI value varies depending on the length of the ITI – with each timestep, there is an increasing belief they are in the ‘Pre’ state, with the current value estimate updating to reflect that. The data cannot be explained by the Cue-Context model, in which ITI value is fixed (Fig. 4g). The modeling suggests that the lack of a significant trend in the remaining two conditions is due to the lower variance in value in the pre-cue period, with an average of 0.28 ± 0.87 and 0.46 ± 1.11 licks (mean ± s.d.) in the 2 second pre-cue period in Conditioning and Cued Reward respectively (versus 1.51 ± 1.52 in Degradation), leading to underpowered analysis.

In summary, the ITI state representation is essential to explaining the relative effects of contingency degradation and additional cued rewards on the Odor A response. Complex ITI representations, such as CSC, are inefficient, whereas modeling it as a single state (Cue-Context), does not capture the heterogeneity of the ITI. Our Belief-State model, representing the ITI using two states, is sufficient to explain the experimental results.

### Additional aspects of dopamine responses and model predictions

Having identified a sufficient model for explaining our contingency degradation results, we next examined how well this model matched additional experimental results. Figure 5a visualizes the value as predicted by the Belief-State model across four conditions tested (Conditioning, including Recovery; Degradation, Cued Reward and Extinction). In the Odor A rewarded trial, the value during the ISI remained unchanged in the first three conditions, and significantly decreased in Extinction, closely mirroring the (prospective) reward probability *P*(*R*|*A*). For the reasons discussed above, the pre-ISI period, reflecting the pre-transition state (‘Pre’), showed a modest increase in the Cued Reward case and a significant rise in the Degradation condition. The TD errors upon Odor A presentation, reflective of the difference in value between Pre state and the first ISI substate, diminished in both Degradation and Extinction. In both these conditions, contingency is reduced by increasing *P*(*R*|*A*−) and decreasing *P*(*R*|*A*), respectively^52^. Notably, our model suggested two distinct mechanisms underlying these two processes: an increase in Pre state value in Degradation and a decrease in ISI value in Extinction (Fig. 5c). Our Belief-State TD learning model matched the experimental results well (Fig. 5b, d), including the Extinction data.

**Figure 5.**
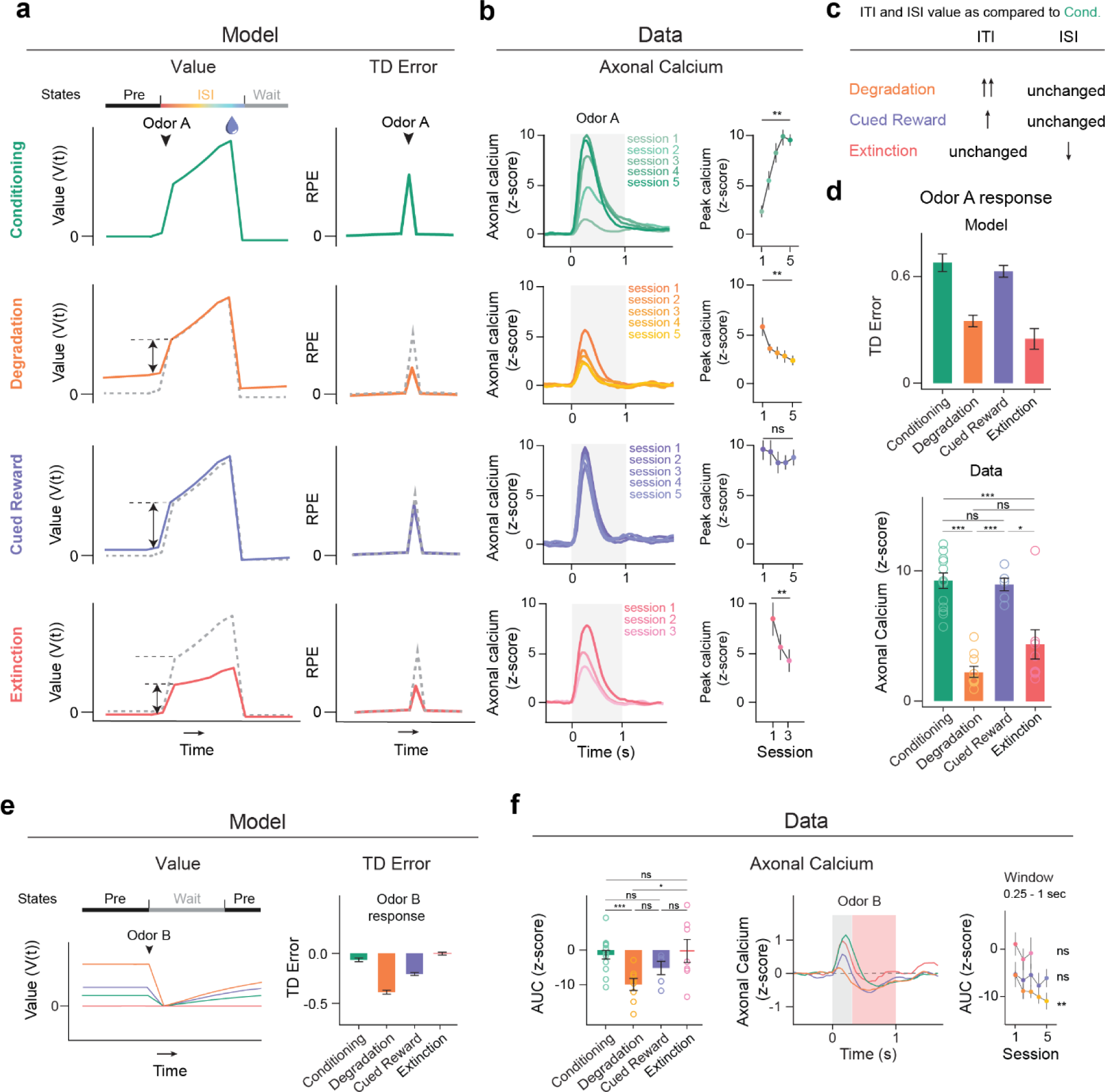
| Belief-State model’s predictions recapitulate additional experimental data. (a) Plots averaged from one representative simulation of Odor A rewarded trial (*n* = 4,000 simulated trials) for four distinct conditions using the Belief-State model. Graphs are for the corresponding value function (left) and TD error (right) of cue response for Odor A rewarded trials. (b) Signals from dopamine axons (mean) across multiple sessions of each condition (left). Mean peak dopamine axonal calcium signal (z-scored) for the first to last session in Phase 2 for four contingency conditions (right). Error bars represent SEM. ns, *P* >0.05; **, *P* < 0.01, Student’s paired *t*-test. The Belief-State model captures the modulation of Odor A dopamine response in all conditions. (c) Degradation, Cued Reward and Extinction conditions differ in how their ITI and ISI values change compared to Conditioning phase. (d) Mean peak TD error by Belief-State model and dopamine axonal signal (z-scored) to Odor A for four distinct conditions. Error bars represent SEM. ns, *P* > 0.05; *, *P* < 0.05; ***, *P* < 0.001, Welch’s *t*-test. The model’s prediction captured well the pattern in the dopamine data. (e) Averaged traces from a representative simulation of Odor B trial (*n* = 4,000 simulated trials) across four distinct conditions using the Belief-State model. Graphs are for the value function and TD errors of cue response for Odor B trials. (f) Z-scored dopamine axonal signals to Odor B quantified from the red shaded area to quantify the later response only. Bar graph (left) shows mean z-scored Odor B AUC from 0.25s-1s response from the last session of each condition. Error bars are SEM. * *P* < 0.05; ***, *P* < 0.001, Welch’s *t*-test. Line graph (right) shows mean z-scored AUC over multiple sessions for each condition. Statistical analysis was performed on data from the first and last sessions of these conditions. Error bars are SEM.

The model predicts another distinct difference between degradation and extinction: degradation affects TD error for all cues due to changes in the shared Pre state value, while extinction impacts only the specific cue undergoing extinction. Accordingly, we examined the Odor B trials. In the Belief-State model, Odor B is a transition from the Pre to Wait state, and thus the TD error is the difference between these two state values. We expected the most negative response in the Deg group, owing to a higher Pre state value, and relatively unchanged ‘Wait’ value. We also expected an unchanged response in Extinction in comparison to Conditioning. Experimentally, the response to Odor B was biphasic, featuring an initial positive response followed by a later negative response. Such a biphasic response has been previously noted, with general agreement that the second phase is correlated with value^53^. By quantifying the later response (250ms-1s window), there was a close match between the model prediction and the data for Odor B responses (Fig. 5e, f).

The Belief-State model shows that TD errors at reward omission are based on the difference between the final ISI substate and Wait state values. The Wait state value, generally lower than the Pre state value, has minor changes across conditions. This results in consistent TD errors at reward omission across Conditioning, Degradation, and Cued Reward conditions due to similar ISI values, but a significant reduction in Extinction due to a lower ISI value, which closely aligned with the experimental results (Extended Data Fig. 6). TD errors at predicted rewards, reflecting the difference between actual reward and ISI values, exhibit minimal changes across Conditioning, Degradation and Cued Reward conditions, which is also consistent with the data.

In total, the above results indicate that the TD model with proper task states can effectively recapitulate nearly all aspects of phasic dopamine responses across various trial types and task events.

### Recurrent neural networks that learn to predict values through TD learning can explain dopamine responses

The models discussed above, while effective, are ‘hand-crafted’ and tuned to our particular task setting. While there is evidence that dopamine neurons rely on belief-state inference in computing TD error^50,51,54^, the question of how the brain learns such a state-space is less well understood. Previous work has shown that RNNs, trained to estimate value directly from observations (‘value-RNNs’), develop belief-like representations despite not being explicitly trained to do so^55^. This approach substitutes hand-crafted states for an RNN that is simply given the same odor and reward observations as the animal (Fig. 6a).

**Figure 6.**
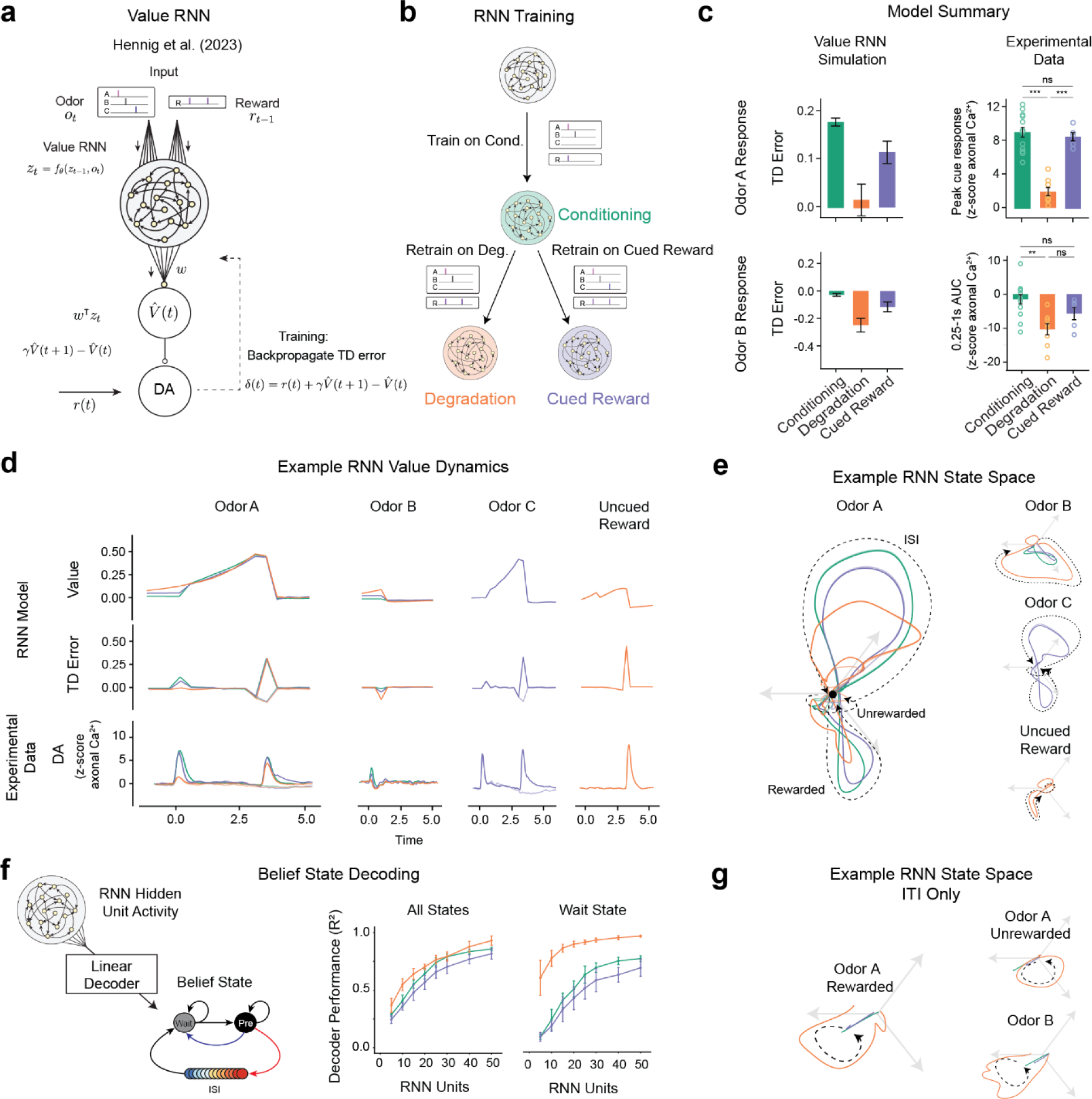
| Value-RNNs recapitulate experimental results using state-spaces akin to hand-crafted Belief-State model. (a) The Value-RNN replaces the hand-crafted state space representation with an RNN that is trained only on the observations of cues and rewards. The TD error is used to train the network. (b) RNNs were initially trained on simulated Conditioning experiments, before being retrained on either Degradation or Cued Reward conditions. (c) The predictions of the RNN models (mean, error bars: SD) closely match the experimental results. (d) Example value, TD error, and corresponding average experimental data from a single RNN simulation. Notably, decreased Odor A response is explained by increased value in the pre-cue period. (e) Hidden neuron activity projected into 3D space using CCA from the same RNNs used in (d). The Odor A ISI representation is similar in each of the three conditions, and similar to the Odor C representation. Odor B representation is significantly changed in the Degradation condition. (f) Correspondence between RNN state space and Belief-State model. A linear decoder was trained to predict beliefs using RNN hidden unit activity. With increasing hidden layer size, the RNN becomes increasingly belief-like. The improved performance of the decoder for the Degradation condition is explained by better decoding of the Wait state. Better Wait state decoding is explained by altered ITI representation: (g) Same RNNs as in (d) and (e), hidden unit activity projected into state-space as (e) for the ITI period only reveals ITI representation is significantly different in the Degradation case.

Here, we applied the same value-RNN to our contingency manipulation experiments. We generated training sets, consisting only of the odor and reward timings, that matched the three conditions. The RNNs were first trained on the Phase 1 Conditioning task and then either on the Phase 2 contingency Degradation or Cued Reward conditions (Fig. 6b). Several RNNs were trained with different numbers of hidden units, from 5 to 50.

The trained RNNs closely matched the experimental results (example 50 unit RNN presented in Fig. 6c). Like the TD models used in the above section, the decrease in Odor A response is explained by an increase in the value during the ITI period, not a shift in the value during the ISI (example Fig. 6d).

We were interested in understanding the inferred state spaces used by the RNN models. To visualize this, we applied canonical correlation analysis (CCA)^56,57^ to align the activity of the hidden units between the RNNs for each condition for all conditions.

In all conditions, without any stimuli, the RNN’s activity will decay to a fixed point (here plotted as the origin, Extended Data – Video 1). This can be understood as the Pre-transition state. In all conditions, the Odor A trajectory is similar, indicating a shared representation of the ISI period (Fig. 6e). Furthermore, in the Cued Reward condition, the Odor C trajectory is nearly identical to that of Odor A, potentially reflecting generalization. In the Degradation condition, delivering Odor B causes a trajectory that is significantly longer than the other two conditions, potentially corresponding to the Wait state.

To compare the state space of the value-RNN to the Belief-State model, we calculated the beliefs at each given time point in the simulated experiment and used a linear regression to relate the hidden unit activity. As previously noted^55^, the unit activity became more belief-like with more hidden units (Fig. 6f). Notably, the regression performance, as quantified by *R*^2^ (see Methods), was higher for the Degradation condition at each hidden layer size. This is explained by better performance on the Wait state (Fig. 6f, right panel). As evident in the visualized activity in the state spaces, the RNNs trained on the Degradation condition developed distinct trajectories in the ITI compared to the other two conditions (Fig. 6g), taking a longer period of time to return to the fixed ITI point and following a similar pattern regardless of the particular trial type. In all RNNs that successfully predicted degradation effect, the Wait state readout had a minimum performance of *R*^2^ = 0.57. This suggests that it is the delivery of rewards during the ITI that reshapes the state space to be heterogeneous, while in the other conditions this is not necessary and thus the ITI has a relatively fixed state space representation. That the RNN can learn a belief-like representation from limited information, using only the TD error as feedback, suggests a generalized method by which the brain can construct state spaces using TD algorithms.

### A retrospective learning model, ANCCR, cannot explain the dopamine responses

While our analysis using the TD learning models with explicit state representations and the value-RNNs suggest that TD learning models are sufficient to explain our experimental results, we have not yet considered whether alternative definitions of contingency would also provide an account of our results. The ANCCR (adjusted net contingency for causal relations) is a recently described new model, proposed as an alternative account of the TD explanation of dopamine activity (Fig. 7a)^31^. The authors have previously shown that this model can account for contingency degradation^31,32^ and suggested that TD accounts could not.

**Figure 7.**
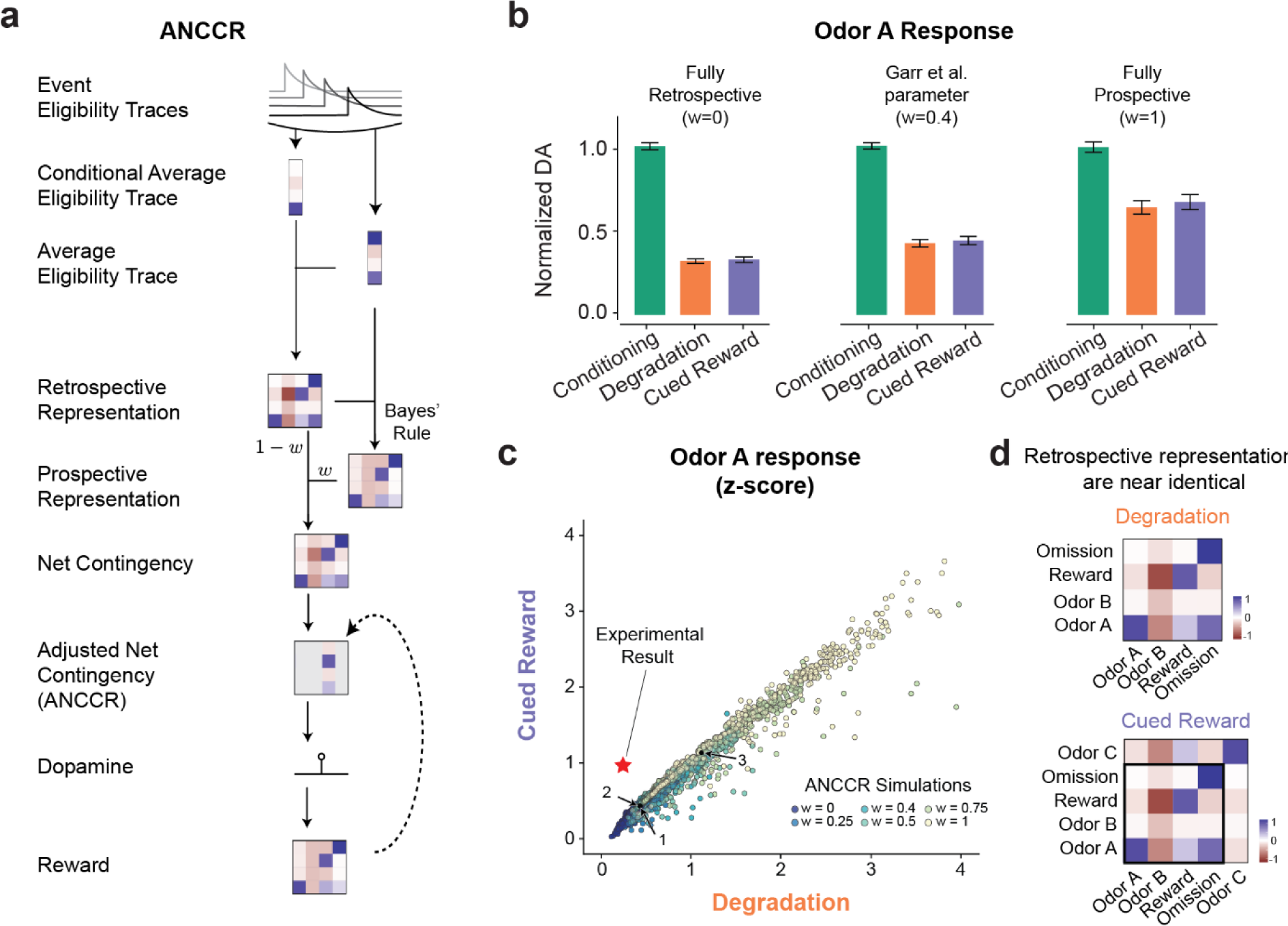
| ANCCR does not explain the experimental results. (a) Simplified representation of ANCCR model. Notably the first step is to estimate retrospective contingency using eligibility traces. (b) Simulations of the same virtual experiments used in Figure 3 using ANCCR, using the parameters in Garr et al., 2023 varying the prospective-retrospective weighting parameter (w). Error bars are SD. In all cases the predicted odor A response is similar in the Degradation and Cued Reward conditions. (c) No parameter combination explains the experimental result. Searching 21,000 parameter combinations across six parameters (T ratio = 0.2-2, α = 0.01-0.3, k = 0.01-1 or 1/(mean inter-reward interval), w =0-1, threshold = 0.1-0.7, αR = 0.1-0.3). Experimental result plotted as a star. Previously used parameters (Garr et al., 2023 as 1, Jeong et al., 2022 as 2 and 3) indicated. Dots are colored by the prospective-retrospective weighting parameter (w), which has a strong effect on the magnitude of Phase 2 response relative to Phase 1. (d) As the contingency is calculated as the first step, and the contingencies are similar in Degradation and Cued Reward conditions, there is little difference in the retrospective contingency representation between the two conditions, explaining why regardless of parameter choice ANCCR predicts similar responses.

ANCCR builds upon the authors’ previous observation that the retrospective information (‘which cues precede reward?’) can be used to explain animal behavior previously unexplained by prospective accounts^33^. Accordingly, the ANCCR model begins with the calculation of the retrospective contingency, using eligibility traces as a principled method to compute contingency in continuous time, rather from trial-by-trial probabilities. At the time of a reward (or a ‘meaningful event’), the difference between the eligibility of cues at the time of reward and the average cue eligibility is computed. This generalizes the trial-based definition of Δ*P*_*retro*_(*A*) to continuous time. From this retrospective contingency and the average event rates, the model proposes the prospective contingencies are inferred using a Bayes-like computation.

From the prospective and retrospective contingencies, a weighted sum (‘net’) contingency is calculated for all pairs of events. This map can be used to calculate the change in expectation of reward for a given event, considering other explanations. It is this ‘adjusted net contingency’ that ANCCR proposes is represented in the dopamine signal.

To test the ANCCR model, we used the authors’ published code to model the same 25 simulated experiments used in our TD modeling. For this experiment, ANCCR has 12 parameters, at least 6 of which have a significant impact on the modeled response. We first tried using the parameters published in Garr et al. (2023) and Jeong et al. (2022); we present results using the Garr parameters because they are closer to our experimental results. While the ANCCR model accurately predicted a decreased response for Odor A during contingency degradation, it predicted a similar response in the Cued Reward condition, conflicting with the experimental results (Fig. 7b). We varied the parameter (*w*) which controls the relative amount of retrospective and prospective information used to calculate the contingency. This parameter controlled the relative size of the decrease, while sensitive to parameter choice governing the eligibility decay rate and learning rates; in general the halving of *P*(*A*|*R*) produces a large decrease in the retrospective contingency, *P*(*A*|*R*) − *P*(*A*), whereas the increase in *P*(*R*) slightly decreases the prospective contingency, *P*(*R*|*A*) − *P*(*R*).

We next considered whether this was a problem of parameter selection, and therefore simulated the first 5 virtual experiments for the parameter search space used in Garr et al. (2023), trying a total of 21,000 parameter combinations, including those in the two previous studies^31,32^ (indicated as 1, 2 and 3). Fig. 7c plots the Odor A dopamine response in the Degradation and Cued Reward case for each of these combinations, normalized by the response during Conditioning. No parameter combination predicted the correct pattern of experimental results, quantitatively or qualitatively (Fig. 7c).

## Discussion

Here we examined behaviors and VS dopamine signals in a Pavlovian contingency degradation paradigm, including a pivotal control. Our results show that dopamine cue responses, like behavioral conditioned responses, were attenuated when stimulus-outcome contingency was degraded by the uncued delivery of additional rewards. Crucially, neither dopamine signals nor conditioned responses were affected in a control condition in which the delivery of additional rewards were cued by a different stimulus despite a similar number of rewards being administered. Our findings not only demonstrate that the above results were not due to satiety, but also provide key insights into possible mechanisms underlying contingency degradation.

Contrary to claims from previous studies^31,32^, our modeling showed that many aspects of dopamine responses can be comprehensively explained by TD learning models, if the model is equipped with proper state representations reflecting the task structure. These TD learning models also readily explained dopamine cue responses in the control condition with cued rewards – results which strongly violated the predictions of a contingency-based retrospective causal learning model (ANCCR) and the Δ*P* definition of contingency. The results indicate that dopamine signals, as well as conditioned responses, primarily reflected the prospective, but not retrospective, stimulus-outcome relations. Rather than discarding the notion of contingency altogether, we propose that these results point toward a novel definition of contingency grounded in the TD learning framework. These results bear significant implications for the theory of associative learning and the nature of dopamine signals, which help resolve some of previously unresolved controversies.

### TD learning model as a model of associative learning

Historically, Pavlovian contingency degradation paradigms have played a pivotal role in the development of animal learning theories^3,16^, yet the exact underlying mechanisms remain to be determined^17,19^. Here we show that the effect of contingency manipulations, both on behavior and dopamine responses, can be explained by TD learning models. As our systematic investigation revealed, the failure of previous efforts to explaining contingency degradation with TD learning models is due to the use of inappropriate state representations, either not considering the ITI period at all, or modeling it in a simplistic way. We show two types of TD learning models that explain the basic behavioral and dopamine results. The first model (Cue-Context model) uses a contextual stimulus as one of the states that continuously exists throughout the task period, which is equivalent to the cue-competition model traditionally considered in the animal learning theory literature^16,17,19^. The second model (Belief-State model) explicitly models the task structure as transitions across the ISI and the two ITI states (“wait” and “pre-transition”), and TD learning operates on beliefs (posterior probabilities) over these discrete states.

In both models, the reduction in dopamine cue responses occurs due to an increase in the value preceding a cue presentation, which decreases the *change* in value (reward expectation) induced by the cue, rather than due to a decrease in the absolute level of the value induced by the cue. This raises the question of why cue-induced anticipatory licking is reduced during contingency degradation. We provide a potential mechanism: the animal distributes anticipatory behavior depending on the relative values across different states.

Our results favor the Belief-State model over the Cue-Context model; both the dopamine and behavioral data were better explained by the Belief-State model. One could argue that it is unclear whether the animal can learn “sophisticated” state representations such as those used in our Belief-State model. In support of a Belief-State TD learning model, our analysis of anticipatory licking indicated that the reward expectation was modulated in a manner intricately linked to different task states: the ISI, wait, and pre-transition states. Furthermore, we show that recurrent neural networks, trained to predict values (value-RNNs), acquired the activity patterns that can be seen as representing beliefs, merely from observations, without explicitly instructed to develop such representations, similar to our previous work using different behavioral tasks^55^. Critically, when trained on contingency degradation sessions, the value-RNNs developed more heterogenous representations of the ITI, capturing the same phenomenon as the Belief-State model.

It has been shown that TD learning models can explain a wealth of phenomena studied in the animal learning theory literature^30,58^. The present study adds to this list Pavlovian contingency degradation – a classic phenomenon long studied in psychology and now in neurobiology. These results indicate that TD learning models provide a foundation with which to understand associative learning while the RNN-based approach provides a principled way to apply TD learning with minimal assumptions about state representations.

### State representations as population activity dynamics

In RL, the “state” is a critical component which represents the set of observable and inferred variables necessary to compute value and policy. The artifice of the state representations used in neurobiological RL modeling has been criticized^59^. For instance, it is implausible to have separate sets of neurons activated sequentially (i.e. CSCs) for separate cues, particularly if they are to completely tile the ITI, as in the CSC with ITI states model^59^. Furthermore, states are often defined within each “trial”; how can states be defined when there are no obvious trial structures^59^? The success of value-RNNs in replicating aspects of dopamine signals and the acquisition of belief-like representations provides two crucial insights into how biological circuits may represent states.

First, the recent successes of RL on complex machine learning tasks, containing many stimuli and often lacking obvious trial structure, suggests that it is possible to achieve high performance with standard RL techniques^22^. A key ingredient lies in the use of neural networks capable of autonomously learning representations appropriate for specific tasks. Our results with value-RNNs agree with this observation. As shown in our previous work^55^ and in the present work, value-RNNs have a stable fixed point (attractor) corresponding to the ITI state (the Pre-transition state in our Belief-State model). The ITI state is thus an emergent property of training to predict reward. Furthermore, different stimuli induce stimulus-specific trajectories in the population activity state space. We found a close correspondence between population dynamics and the hand-crafted states assumed in Belief-State TD learning models. These results indicate that the population activity patterns in a network, including attractors and stimulus-specific trajectories, represent distinct states such as those assumed in our Belief-State TD learning models. Although the activity representing different trajectories likely involves overlapping sets of neurons, they can be trained to compute values properly by adjusting downstream synaptic weights (as long as the activity patterns for different states are discriminable).

Second, while TD learning models with hand-crafted state representations help develop conceptual understanding, the RNN-based approach can provide insights into how hand-crafted state representations could be implemented in neural networks. In the future, it is of great interest to examine whether neural activity in the brain exhibits patterns of activity predicted by the value-RNN models. The prefrontal cortex is a strong candidate area, receiving dopaminergic innervation from the VTA necessary for appropriate adaptation to contingency degradation in instrumental conditioning^60^. However other areas, such as the hippocampus, also contribute task-relevant information during degradation to the prefrontal cortex^61^ and neural network modeling approaches that reflect the brain’s functional organization (e.g. ^62^) might provide more insight than our model which treats the state-machinery of the brain as a single recurrent neural network.

### Limitations of the ANCCR model as a model of associative learning and dopamine

The present study unveiled limitations of the recently proposed causal learning model, ANCCR^31,32^. The Degradation and Cued Reward conditions are minimally different and thus provide a strong test of the algorithm design. Our results indicate that the ANCCR model fails to explain the observed results despite our extensive examination of its parameter space. Crucially, the ANCCR model suffers from the same flaw as the Δ*P* definition of contingency. While extending the definition to continuous time and considering multiple cues, ANCCR still calculates contingency by subtracting the background event rate, losing information, precluding it from attributing increased value to the background in the same manner as the TD models. Given the similarity in event rate between the conditions, the retrospective representations (and average eligibility trace) remain similar, with the computed retrospective contingencies in the Degradation condition being a subset of the Cued Reward contingencies (Fig 7d). This explains why ANCCR predictions are similar for the two conditions independent of the parameter choice, as the rest of the model depends on this computed retrospective contingency as input. Thus, the failure of the ANCCR to explain the Cued Reward condition reflects the fundamental construction of the ANCCR model.

The failure of the ANCCR model here does not exclude some of the interesting ideas integrated into ANCCR, including how it uses retrospective information to learn the state space. Rather, it is its reliance on contingency that constitutes its core deficit. Other theoretical work has considered how TD algorithms that consider retrospective information may enhance learning performance without explicitly invoking contingency. A recent report^32^ demonstrated that ANCCR is able to explain the dopamine response in outcome-selective contingency degradation. This is a result of the multidimensional tracking of cue-outcome contingencies in ANCCR. We show that both the Belief-State model and the value-RNN, if trained on each reward separately and with total value taken to be the absolute difference of the two separate values, successfully predicts the experimental results of Garr et al. (2023) (Extended Data Fig. 7). A similar approach using “multi-threaded predictive models” was used to successfully explain dopamine data in a different multi-outcome task^44^. While this proposal leaves open questions about how such abstract state representation is implemented biologically (the same being true for ANCCR), it does demonstrate that more complex contingency manipulations can still be explained by TD models. In fact, recent studies have provided evidence for heterogeneous responses to different types of rewards in dopamine neurons^63–65^. While further evidence is required to solidify this understanding, the provisional assumption of multiple value channels shows how TD models for multiple outcomes can potentially be achieved in neural circuitry by concurrently running parallel circuits.

### TD error, contingency and causal inference

Learning predictive or causal relationships requires properly assigning credits to those events that are responsible for the outcomes observed. A key to this process is considering counterfactuals^66^ – would a particular outcome occur had I not seen that cue, or had I not taken that action? In the present study, we show that TD learning models with ITI representations learn and predict value in the time before cue presentation. The cue-associated TD error is then calculated as the difference in value in the presence and absence of that cue. Consequently, computation of TD errors effectively subtracts the prediction of value in the absence of the cue – i.e. the counterfactual prediction. More generally, the computation of TD error or its variants can be seen as subtracting out counterfactuals. In a class of RL algorithms commonly used in artificial intelligence applications (advantage actor-critic algorithms), the actor decides which action to take for a given state and the critic evaluates the action by computing the advantage function, defined as:

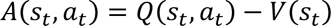

where *Q*(*s*_*t*_, *a*_*t*_) is the state-action value function^67,68^. If this is taken to be the immediate reward of the action plus the expected return of the new state, *Q*(*s*_*t*_, *a*_*t*_) = *r*_*t*_ + *γV*(*s*_*t*+1_) then the advantage function can be approximated by the TD error *A*(*s*_*t*_, *a*_*t*_) = *Q*(*s*_*t*_, *a*_*t*_) − *V*(*s*_*t*_) = (*r*_*t*_ + *γV*(*s*_*t*+1_)) − *V*(*s*_*t*_) = *E*[*δ*_*t*_] (ref. ^69,70^).

As discussed in recent work^70^, in fully observable environments without confounds, the advantage function is exactly equivalent to the Neyman-Rubin definition of causal effect of an action: the difference in outcomes given an action versus outcomes otherwise. In this context, the definitions of causality, contingency and TD error align – all emphasizing the consideration of counterfactual prediction: that is, the difference between potential future outcomes (following action) and the alternative when the action is not taken. TD error can therefore be both a measure of contingency and useful in establishing causal relationships, without invoking retrospective computations.

TD errors improve over the ANCCR and Δ*P* definitions because the comparison to the reward probability of US given CS is not simply the reward probability given absence of CS, but to *V*(*s*), which is the γ-discounted sum of all future rewards given the current state, with the state encapsulating all environmental information. As demonstrated by our modeling, the heterogeneous state representation during the absence of events (the ITI) is critical to the accuracy of our models to match the experimental data.

While these relationships between TD error and contingency hold in fully observable environment, our value-RNN approach may extend these results to more complex/realistic environments. Veitch et al. (2019) has demonstrated network embeddings, like our value-RNN, can reduce the problem of inferring causality to a problem of predicting outcomes^71^. These networks do not require full knowledge of the environment to succeed but rather learn to extract sufficient information to establish causality. Ultimately, TD error could provide pivotal signals for contingency – the essential quantity for causal inference.

## Conclusions

Our results indicate that TD learning models can explain contingency degradation – a phenomenon that was thought to be difficult to explain based on TD learning^31,32,72^. The Belief-State TD model that we used here is “model-free” in the sense that the values are “cached” to each state based on direct experiences, although these states reflect the animal’s knowledge of the transition structure between states which can be regarded as a “world model”^41,43,73^. This suggests that the distinction between “model-free” and “model-based” mechanisms is not as hard-lined as often assumed. The sensitivity to contingency degradation in instrumental behaviors has been used to support the behavior being goal-directed or model-based. Yet, the same type of Belief-State TD model can, in principle, be applied to explain such an effect. In any case, further biological investigations will be needed to constrain mechanisms linking behavior and contingency – the critical variable thought to underlie learning predictive and/or causal relationships. The experimental results and models presented in this study would aid such efforts.

## Methods

### Animals

A total of 31 mice were used. 18 wildtype mice (8 males and 10 females) at 3-6 months of age were used to collect only behavioral data. For fiber photometry experiments, 13 double transgenic mice resulting from the crossing of DAT-Cre (Slc6a3tm1.1(cre)Bkmn; Jackson Laboratory, 006660)^37^ with Ai148D (B6.Cg-Igs7tm148.1(tetO-GCaMP6f,CAG-tTA2)Hze/J; Jackson Laboratory, 030328)^38^ (DAT::cre x Ai148, 7 males and 6 females) at 3–6 months of age were used. Mice were housed on a 12 hr /12 hr dark/light cycle. Ambient temperature was kept at 75 ± 5 °F and humidity below 50%. All procedures were performed in accordance with the National Institutes of Health Guide for the Care and Use of Laboratory Animals and approved by the Harvard Animal Care and Use Committee.

### Surgery

Mice used for fiber photometry recordings underwent a single surgery to implant a multifiber cannula and a head fixation plate 2-3 weeks prior to the beginning of the behavioral experiment. All surgeries were performed under aseptic conditions. Briefly, mice were anesthetized with an intraperitoneal injection of a mixture of xylazine (10 mg/kg) and ketamine (80 mg/kg) and placed in a stereotaxic apparatus in a flat skull position. During surgery, the bone above the Ventral Striatum area was removed using a high-speed drill. A custom multifiber cannula (6 fibers, 200 µm core diameter, 0.37 NA, Doric Lenses) was lowered over the course of 10 min to target 6 subregions in the Ventral Striatum. The regions’ coordinates relative to Bregma (in mm) were: Lateral nucleus accumbens (lNAc, AP:1.42, ML:1.5, DV:-4.5); Medial NAc (mNAc, AP:1.42, ML:1, DV:-4.5); anterior lateral olfactory tubercle (alOT, AP:1.62, ML:1.3, DV:-4.8); posterior lateral OT (plOT, AP:1.00, ML:1.3, DV:-5.0); anterior medial OT (amOT, AP:1.62, ML:0.8, DV:-4.8); posterior medial OT (pmOT, AP:1.00, ML:0.8, DV:-5.0). Dental cement (MetaBond, Parkell) was then used to secure the implant and custom headplate and to cover the skull. Mice were singly housed after surgery and post-operative analgesia was administered for 3 days (buprenorphine ER-LAB 0.5 mg/ml). Mice used for behavioral training underwent a similar surgical process, but only a head fixation plate was implanted.

### Behavioral training

After recovery from headplate-implantation surgery, animals were given ad libitum access to food and water for 1 week. Before experiments and throughout the duration of the experiments, mice were water restricted to reach 85–90% of their initial body weight and provided approximately 1–1.5 mL water per day in order to maintain the desired weight and were handled every day. Mice were habituated to head fixation and drinking from a waterspout 2-3 days prior to the first training session. All tasks were run on a custom-designed head-fixed behavior set-up, with software written in MATLAB and hardware control achieved using a BPod state machine (1027, Sanworks). A mouse lickometer (1020, Sanworks) was used to measure licking as infra-red beam breaks. The water valve (LHDA1233115H, The Lee Company) was calibrated, and a custom-made olfactometer was used for odor delivery. The odor valves (LHDA1221111H, The Lee Company) were controlled by a valve driver module (1015, Sanworks) and a valve mount manifold (LFMX0510528B, The Lee Company). All components were controlled through the Bpod state machine. Odors (1-hexanol, d-limonene, and ethyl butyrate, Sigma-Aldrich) were diluted in mineral oil (Sigma-Aldrich) 1:10, and 30 µL of each diluted odor was placed on a syringe filter (2.7-µm pore size, 6823-1327, GE Healthcare). Odorized air was further diluted with filtered air by 1:8 to produce a 1 liter/min total flow rate. The identity of the rewarded and non-rewarded odors were randomized for each animal.

In Conditioning sessions, there are three types of trials: (1) trials of Odor A (40% of all trials) associated with a 75% chance of water delivery after a fixed delay (2.5 s), (2) trials of unrewarded Odor B (20% of all trials) as control to ensure that the animals learned the task, and (3) background trials (40% of all trials) without odor presentation. Rewarded odor A trials consists of 2s pre-cue period, 1s Odor A presentation, 2.5s fixed delay prior to a 9 µL water reward and 8s post-reward period. Unrewarded Odor B trials consist of a 2s pre-cue period, 1s Odor B presentation, and 10.5s post-odor period. Background trials in the Conditioning phase span a 13.5s eventless period. Trial type was drawn pseudo-randomly from a scrambled array of trial types maintaining a constant trial type proportion. Inter-trial-intervals (ITI) following the post-reward period were drawn from an exponential distribution (mean: 2s).

Learning was assessed principally by anticipatory licking detected at the waterspout for each trial type, with mice performing 100-160 trials per session until they reach an asymptotic task performance, typically after 5 sessions.

After the Conditioning phase, the mice were divided into three groups to undergo different conditions: Degradation (Deg group), Cued Reward (CuedRew group), and Conditioning (Cond group). The Deg group experienced contingency decrease during the Degradation phase. In the Degradation phase, Odor A still delivers water reward with 75% probability, and Odor B remains unrewarded. The difference was the introduction of uncued rewards (9 µL water) in 75% of background trials to diminish the contingency. Animals underwent 5 sessions, each with 100-160 trials, to adapt their conditioned and neural responses to the new contingency. Degradation changed the cue value relative to the background trial but did not impact the reward identity, reward magnitude, or delay to/probability of expected reward.

The CuedRew group was included to account for potential satiety effects due to the extra rewards the Deg group mice received in the background trials. Unlike the Deg group, the CuedRew group’s background trials were substituted with rewarded Odor C trials, where mice received additional rewards signaled by a distinct odor (Odor C). Rewarded odor C trials have the same trial structure as the rewarded odor A trials and animals were given 5 sessions, with 100-160 trials each, to adapt their conditioned response and neural responses to this manipulation.

The Cond group proceeded with an additional five Conditioning sessions, keeping the trial structure and parameters unchanged as in the Conditioning phase.

Post-degradation: eight mice were randomly chosen from the Deg group for the reinstatement phase, replicating the initial Conditioning conditions. After three reinstatement sessions, once the animals’ performance rebounded to pre-degradation levels, we initiated the extinction process. This involved the delivery of both odors A and B without rewards, effectively extinguishing the cue-reward pairing. To mitigate the likelihood of animals generating a new state to account for the sudden reward absence, a shorter reinstatement session was conducted prior to the Extinction session on the extinction day. Extinction was conducted over three days, each day featuring 100-160 trials. After Extinction, a second reinstatement session was implemented, re-introducing the 75% reward contingency for odor A. All eight animals resumed anticipatory licking within ten trials during this reinstatement.

### Fiber photometry

Fiber photometry allows for recording of the activity of genetically defined neural populations in mice by expressing a genetically encoded calcium indicator and chronically implanting optic fiber(s). The fiber photometry experiment was performed using a bundle-imaging fiber photometry setup (BFMC6_LED(410-420)_LED(460-490)_CAM(500-550)_LED(555-570)_CAM(580-680)_FC, Doric Lenses) that collected the fluorescence from a flexible optic fiber bundle (HDP(19)_200/245/LWMJ-0.37_2.0m_FCM-HDC(19), Doric Lenses) connected to a custom multifiber cannula containing 6 fibers with 200-μm core diameter implanted during surgery. This system allowed chronic, stable, minimally disruptive access to deep brain regions by imaging the top of the patch cord fiber bundle that was attached to the implant. Interleaved delivery 473 nm excitation light and 405 nm isosbestic light (using LEDs from Doric Lenses) allows for independent collection of calcium-bound and calcium-free GCaMP fluorescence emission in two CMOS cameras. The effective acquisition rate for GCaMP and isosbestic emissions was 20Hz. The signal was recorded during each session when the animals were performing the task. Recording sites which had weak or no viral expression or signal were excluded from analysis.

The global change of signals within a session was corrected by a linear fitting of dopamine signals (473nm channel) using signals in the isosbestic channel during ITI and subtracting the fitted line from dopamine signals in the whole session. The baseline activity for each trial (F_0 each_) was calculated by averaging activity in the pre-stimulus period between −2 to 0 seconds before an odor onset for odor trials or water onset for uncued reward trials. Z-score was calculated as (F − F_0 each_)/STD_ITI with STD_ITI the standard deviation of the signal during the ITI.

To quantify Odor A responses, we looked for ‘peak responses’ by finding the point with the maximum absolute value during the 1-s window following the stimulus onset in each trial. To quantify Odor B responses, we measured area under curve by summing the value during the 250 ms to 1s window following the stimulus onset in each trial. This is to separate out the initial activation (odor response) that we consistently observed, and which may carry salience or surprise information independent of value. To quantify reward responses, we looked for ‘peak responses’ by finding the point with the maximum absolute value during the 1-s window following the reward onset in each trial. To quantify reward omission responses, we looked for area under curve by summing the value during the 0-1.5s window following the reward omission in each trial.

### Histology

To verify the optical fiber placement and GCaMP expression, mice were deeply anesthetized with an overdose of ketamine-medetomidine, and perfused transcardially with 0.9% saline followed by 4% paraformaldehyde (PFA) in PBS at the end of all experiments. Brains were removed from the skull and stored in PFA overnight followed by 0.9% saline for 48 hours. Coronal sections were cut using a vibratome (Leica VT1000S). Brain sections were imaged using fluorescent microscopy (AxioScan slide scanner, Zeiss) to confirm GCaMP expression and the location of fiber tips. Brain section images were matched and overlaid with the Paxinos and Franklin Mouse Brain Atlas cross-sections to identify imaging location.

### Computational Modeling

#### Simulated Experiments

To compare the various models, we generated 25 simulated experiments of Cond, Deg and CuedRew groups, matching trial statistics to the experimental settings, but increasing the number of trials to 4,000 in each phase to allow to test for steady-state response in both these TD simulations and the ANCCR simulations. We then calculated the state representation of the simulated experiments for each of four state representations (CSC with and without ITI states, Context-TD, Belief-State model, detailed below) and ran the TD learning algorithm with no eligibility trace, called TD(0), using these state representations (Fig. 3a). While TD(0) has a learning rate parameter (*α*), it did not influence the steady-state results, which are presented, and thus the only parameter which influenced the result was *γ*, the temporal discount factor, set to 0.925 for all simulations using a timestep of Δt = 0.2s (Extended Data Fig. 4 shows the *γ* parameter search space). Code for generating the simulated experiments and implementing the simulations can be found at: https://github.com/mhburrell/Qian-Burrell-2024

##### CSC-TD model with and without ITI states

We initially simulated the Conditioning, Degradation, and Cued Reward experimental conditions using the CSC-TD model, adapted from Schultz et al.^25^. The cue length was fixed at 1 unit of time, with time unit size set to 0.2 s, and the ISI was matched to experimental parameters at 3.5 s. Simulated cue and reward frequencies were matched to experimental parameters, separately simulating Conditioning then Degradation and Conditioning then Cued Reward. In complete serial compound, also known as tapped-delay line, each cue results in a cascade of discrete substates that completely tile the ISI. TD error and value were then modelled using a standard TD(0) implementation^21^, using *α* = 0.1, *γ* = 0.925. Reported values are the average of the last 200 instances averaged for 25 simulations. The model was run with states tiling the ISI only (CSC) or tiling the ISI and ITI until the next cue presentation (CSC with ITI states).

##### Context-TD model

The Context-TD model, which is an extension of the CSC-TD model, includes context as an additional cue, but otherwise identical to the CSC simulations. For each phase (Conditioning, Degradation, Cued Reward) a separate context state was active for the entire phase, including the ISI and ITI. This corresponds to the additive Cue-Context model previously described^16,17,19^. TD errors reported are the average of the last 200 instances averaged for 25 simulations, except for Extinction which corresponded to third day of training.

##### Belief-State model

We simulated the TD error signaling in all four conditions (Conditioning, Degradation, Cued Reward, Extinction) using a previously described belief-state TD model^50^. For comparison to the CSC based models described above, we had a total of 19 states, 17 capturing the ISI substates (3.5s in 0.2s increments, as in the CSC model). State 18 we termed the ‘Wait’ state and state 19 the ‘pre-transition’ or ‘pre’ state. In the Belief-State model it is assumed the animal has learned a state transition distribution. We computed the transition matrix by labelling the simulated experiments with state, labelling the fixed post-US period as the Wait state and the variable ITI as the Pre state and then empirically calculating the transition matrix for that simulation. While the post-US and variable ITI periods were used to estimate the rate of transition between the Wait and Pre states, because we assumed a fixed probability of transition, these should not be considered identical – rather the implicit assumption in modeling with a fixed probability is that the time in the Wait state is a geometric random variable.

The belief-state model also assumes that the animal has learned a probability of distributions given the current state, encoded in an observation matrix. In our implementation there are five possible observations: Odor A, B, C, reward and null (no event). Like the transition matrix, the observation matrix was calculated empirically from the simulated experiments. Fig 3b represents schematically the state-space of the Belief-State model:Odor A (and C in Cued Reward) are observed when transitioning from Pre to the first ISI state; reward is observed in transition from the last ISI state to Wait, Odor B (and reward in Deg) are observed when transitioning from Pre to Wait. We did not consider how the details of how the transition and observation matrices may be learnt on a trial-by-trial basis as the steady-state TD errors are not dependent on this implementation. As for the other models, the TD errors reported are the average of the last 200 instances averaged over 25 simulations, except for Extinction which corresponded to the third day of training.

A relative value metric was used as a potential explanation of the decrease in licking during the ISI in the Degradation condition (Extended Data Fig 5). Relative value at time t was computed as value at time t (as defined and simulated by the Belief-State TD model) divided by the total value of the entire session, multiplied by the total number of rewards in a session.

### RNN Modeling

We implemented value-RNNs, as described previously^55^, to model the responses in the three conditions (Conditioning, Degradation, Cued Reward). Briefly, simulated tasks were generated to match experimental parameters using a time step of 0.5s. We then trained recurrent network models, in PyTorch, to estimate value. Each value-RNN consisted of between 5 and 50 GRU cells, followed by a linear readout of value. The hidden unit activity, taken to be the RNN’s state representation, can be written as *z*_*t*_ = *f*_*ϕ*_(*o*_*t*_, *z*_*t*−1_) given parameters ϕ. The RNN’s output was the value estimate *V*_*t*_ = *w*^⊤^*z*_*t*_ + *w*_0_, for *z*_*t*_, *w* ∈ ℝ^*H*^ (where H is the number of hidden units) and *V*_*t*_, *w*_0_ ∈ ℝ. The full parameter vector *θ* = [*ϕ w w*_0_]was learned using TD learning. This involved backpropagating the gradient of the squared error loss 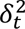 = (*r*_*t*_ + *γV*_*t*+1_ – *V*_*t*_)^2^ with respect to *V*_*t*_ on episodes composed of 20 concatenated trials. The timestep size was 0.5 s and γ was 0.83 to match the 0.925 for 0.2 s timesteps used in the TD simulations, such that both had a discount rate of 0.67 per second.

Prior t o training, the weights and biases were initialized with the PyTorch default. To replicate the actual training process, we initially trained the RNNs on the Cond simulations, then on either the Degradation or Cued Reward conditions (Fig 6b). Training on the Cond simulations for 300 epochs on a session of 10,000 trials, with a batch size of 12 episodes. Parameter updates used Adam with an initial learning rate of 0.001. To replicate the actual training process, we initially trained the RNNs on the Cond simulations, then on either the Degradation or Cued Reward conditions (Fig 6b). To simulate animals’ internal timing uncertainty, the reward timing was jittered 0.5 seconds on a random selection of trials. The model summary plots (Fig 6c, Extended Data Fig 6) presents mean RPE for each event. Exemplar trials shown in Fig 6 have the jitter removed for display purposes.

To visualize the state space used, we performed a two-step canonical correlation analysis process adapting methods used to identify long-term representation stability in the cortex^74^. Briefly, in each condition, we applied principal component analysis (PCA) to identify the principal components (PCs) that explained 80% of the variance (mean number of components = 4.26), then used CCA (Python package pyrcca) to project the PCs into a single space for all conditions. CCA finds linear combinations of each of the PCs that maximally correlate – allowing us to identify hidden units encoding the same information in the different RNNs. We then used the combination of PCA and CCA to create a map from hidden unit activity to a common state.

We measured belief *R*^2^ as previously described^55^. For each simulation, we calculated the beliefs from the observations of cues and rewards. We then used multivariate linear regression to decode these beliefs from hidden unit activity. To evaluate model fit, calculate the total variance explained as: *R*^2^ = 1 − 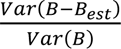, where *B_est_* is the estimate from the regression and *Var*(*X*) = 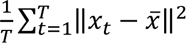

### ANCCR model

The ANCCR model is a recent alternative explanation of dopamine function^31^. While two previous studies have tested contingency degradation with ANCCR, they did not include the cued-reward controls. We implemented the ANCCR model using the code provided on the repository site (https://github.com/namboodirilab/ANCCR) and matching the simulation parameters to the experiment. We used the set of parameter values used in the previous studies, trying both Jeong et al., (2022) and Garr et al. (2023). The total parameter space searched was: T ratio = 0.2-2, α = 0.01-0.3, k = 0.01-1 or 1/(mean inter-reward interval), w =0-1, threshold = 0.1-0.7, αR = 0.1-0.3. The presented results use the parameters from Garr et al. (2023), as they were a better fit (T ratio = 1, α = 0.2, k = 0.01, w = 0.4, threshold = 0.7, αR = 0.1). Additionally, we varied the weight of prospective and retrospective processes (w) to examine whether the data can be explained better by choosing a specific weight. Data presented are the last 200 instances averaged for the same 25 simulations used in the TD simulations.

### Outcome Specific Degradation Modeling

To model outcome-specific degradation we adapted both our Belief-State model and RNN models. For the Belief-State, we estimated the transition and observation matrix for the experiments described in Garr et al., 2023 (depicted in Extended Data Fig 8a) as described for our experiment, using a time step of 1s. As there were two rewarded trial types, we had representations of two ISI periods (termed ISI 1 and ISI 2, depicted in Extended Data Fig 8). The model was initially trained on the liquid reward (setting r =1 when observing liquid reward, r=0 when observing food reward) and the average TD error calculated for each trial type. We then trained on only the food reward. The total TD error was calculated as the absolute difference between the TD error on each reward type.

For the RNN models, we similarly adjusted the timestep to 1s and trained on simulated experiments to match the experimental parameters. Rather than training separately, the model was trained on both simultaneously, training to produce an estimate of the value of the liquid reward and an estimate of the food reward, then using the 2-dimensional vector TD error to train the model. This ensures a single state space is used to solve for both reward types. Total TD error was calculated as the absolute difference on each reward type post-hoc.

### Statistical analysis

Data analysis was performed using third party packages (e.g. Scipy, Statsmodel, etc.) in Python. All code used for analysis is available on request. Our behavioral data and dopamine response data have passed the normality test. For statistical comparisons of the mean, we used Student’s *t*-test with a significance threshold of 0.05, adjusted with the Bonferroni correction. We used Welch’s test for dopamine response to various events due to inequal variance between groups. Paired *t*-tests were conducted when the same mouse’s performance was being compared across two different sessions. No statistical methods were used to predetermine sample sizes, but our sample sizes are similar to those reported in previous publications. The assumptions of the t-test were tested using the Shapiro-Wilk test to check for normality and Levene’s test to check for equal variance.

## Material availability

### Data availability

The behavioral and fluorometry data will be shared at a public deposit source.

### Code availability

The model code will be attached as Supplementary Data. All other conventional codes used to obtain the results will be available from a public deposit source.

## Acknowledgements

We thank Hao Wu and Nune Martiros for technical assistance on the behavioral code design; Mitsuko Uchida for discussion and advice on task design; Catherine Dulac, Florian Engert, and all lab members from Naoshige Uchida lab and Venkatesh Murthy lab for discussion. This work was supported by grants from the National Institute of Health (U19 NS113201, R01DC017311), the Simons Collaboration on Global Brain, the Air Force Office of Scientific Research (FA9550-20-1-0413), the Human Frontier Science Program (LT000801/2018 to S.M.), the Harvard Brain Science Initiative, and the Brain and Behavior Research Foundation (NARSAD Young Investigator no. 30035 to S.M.).

## Author information

These authors contributed equally: Lechen Qian, Mark Burrell

## Contributions

L.Q., N.U., and V.N.M conceived the conceptual framework and designed the behavioral tasks and recording experiments. L.Q. conducted all experiments and data analysis. S.M. established the multifiber photometry system and supplied the transgenic mice. M.B., N.U. and L.Q. discussed the modeling framework. M.B. constructed all the TD learning models and conducted the analysis. J.H. constructed RNN models. The RNN modeling results were analyzed by M.B., J.H. and L.Q. The results were discussed and interpreted by L.Q., N.U., M.B., J.H, S.G. and V.N.M. The manuscript was written by L.Q., M.B., and N.U. and all other authors provided feedback.

---

**Extended Data Fig. 1.**
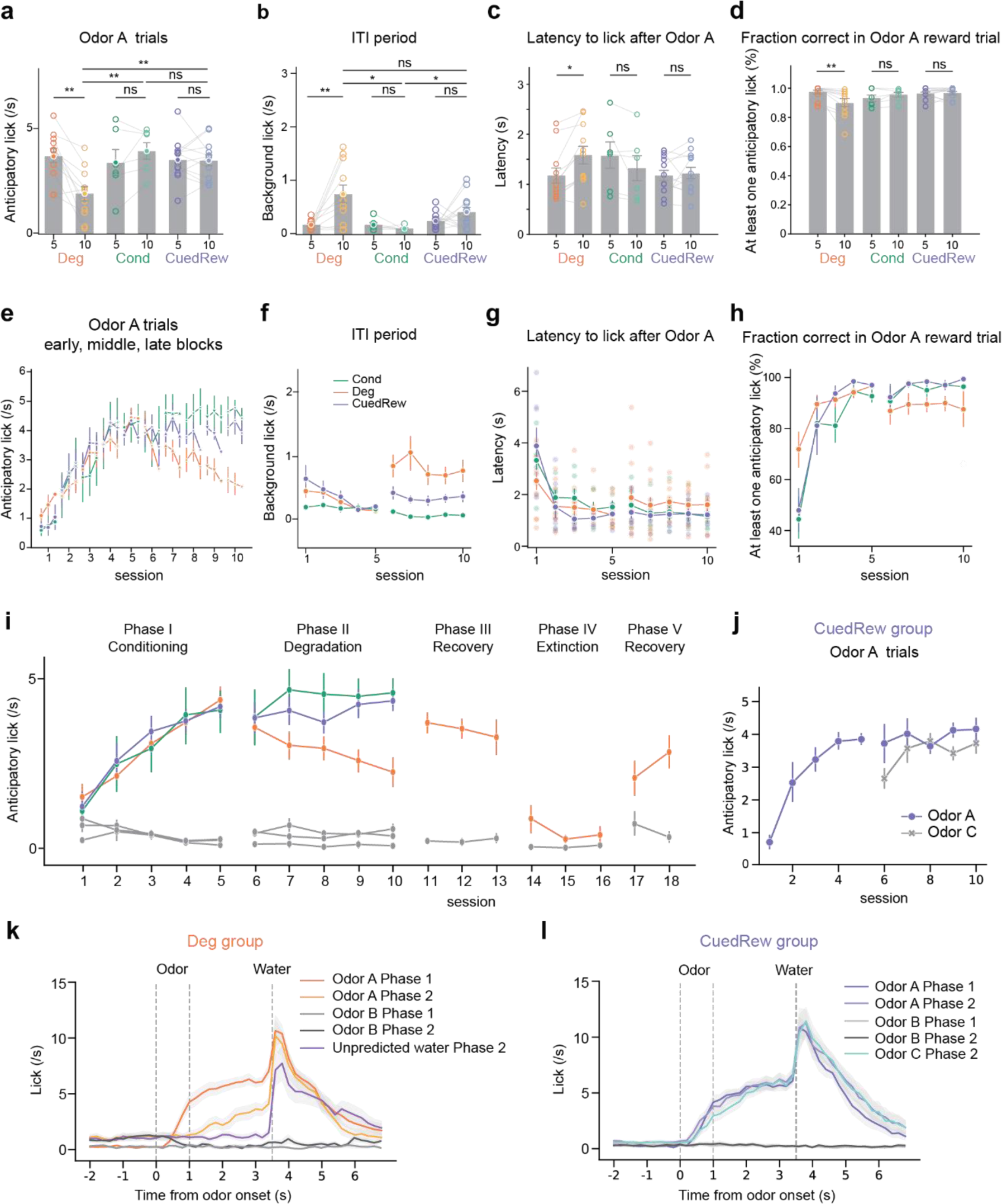
| Population Average Behavior per session. (a, b, c, d) Bar graphs comparing the average number of licks to Odor A during the first 3s post-stimulus (a) and during ITI (b), latency to lick (c), and fraction correct (d) in the final sessions of phase 1 and phase 2 for Deg, Cond, and CuedRew groups. Error bars represent SEM. Asterisks denote statistical significance: ns p > 0.05, **p < 0.01, paired Student’s t-test (e) Session-wise variation in anticipatory licking for Odor A trials, broken down into early, middle, and late blocks, for all groups. (f, g, h). Line graphs showing the average number of licks to Odor A (colored) during ITI (g), latency to lick after Odor A and fraction correct in Odor A trials for each session in the Conditioning, Degradation, and Cued Reward phase (Deg group – orange, n = 11; Cond group – green, n = 6; CuedRew – purple, n=12 mice). (i) Anticipatory licking rate in Odor A trials (colored) and in Odor B trials (grey) across multiple phases: Conditioning (Phase I), Degradation (Phase II), Recovery (Phase III), Extinction (Phase IV), and post-Extinction Recovery (Phase V). (j) Anticipatory licking to Odor C develops quickly compared to Odor A, potentially reflecting generalization. (k, l) PSTH showing the average licking response of mice in Deg group (k) and CuedRew group (l) to the various events. The response is time-locked to the odor presentation (time 0). The shaded area indicates the standard error of the mean (SEM).

**Extended Data Fig. 2.**
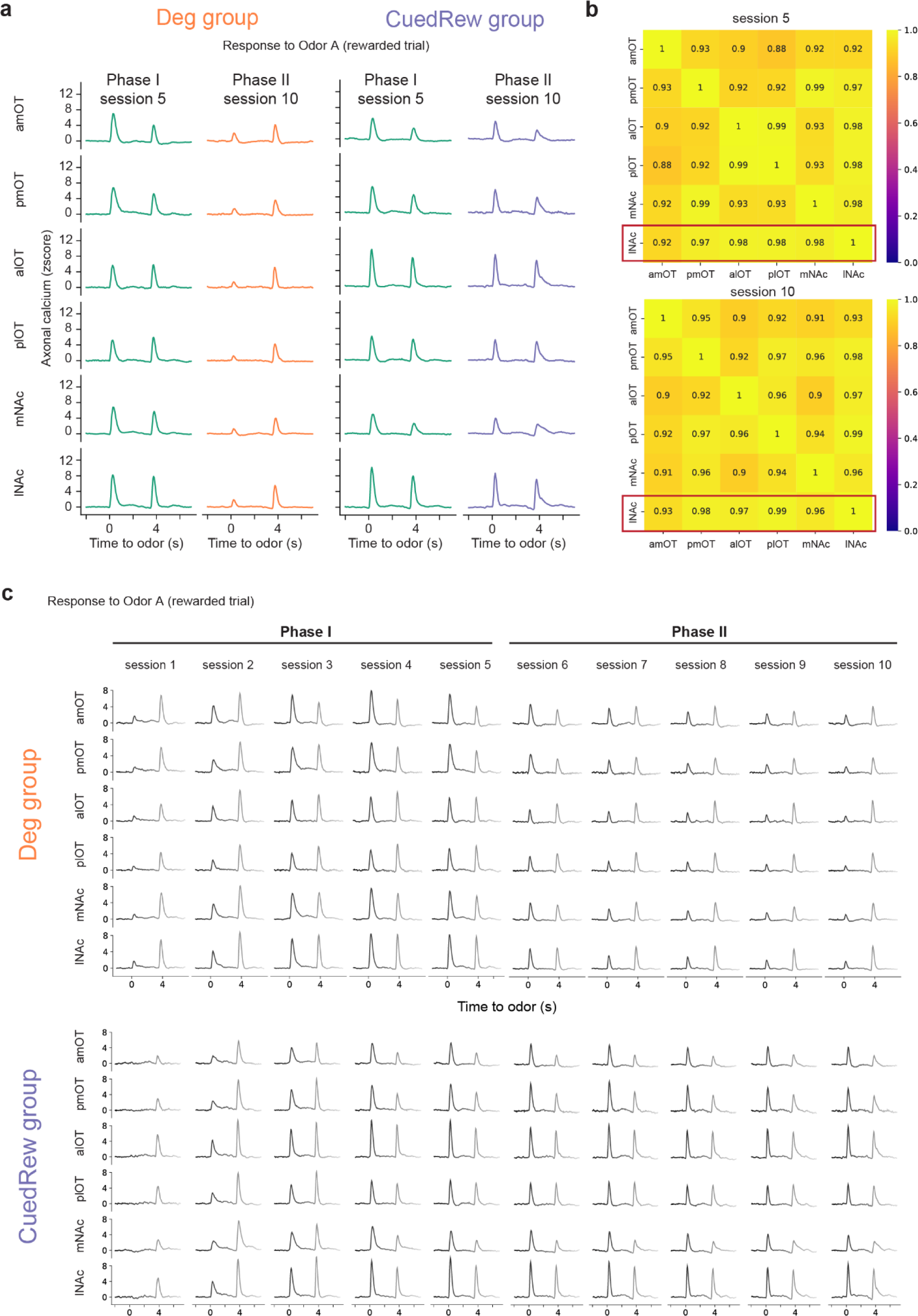
| Dopamine responses are highly correlated across recording sites. (a) Averaged dopamine axonal responses to Odor A during rewarded trials for both Deg group and CuedRew group, depicted for Phase I session 5 and Phase II session 10 across all recorded sites. (b) Correlation matrix for averaged dopamine responses to Odor A during rewarded trials, comparing across sites from the Deg groups during sessions 5 and 10. Cosine similarity was calculated by averaging z-scored responses across trials within animals, then across animals and then computing the cosine similarity between each recording site. (c) Population average dopamine responses to Odor A in rewarded trials across sessions 1 to 10 for both Deg and CuedRew groups, detailing the changes in response through Phase I and Phase II.

**Extended Data Fig. 3.**
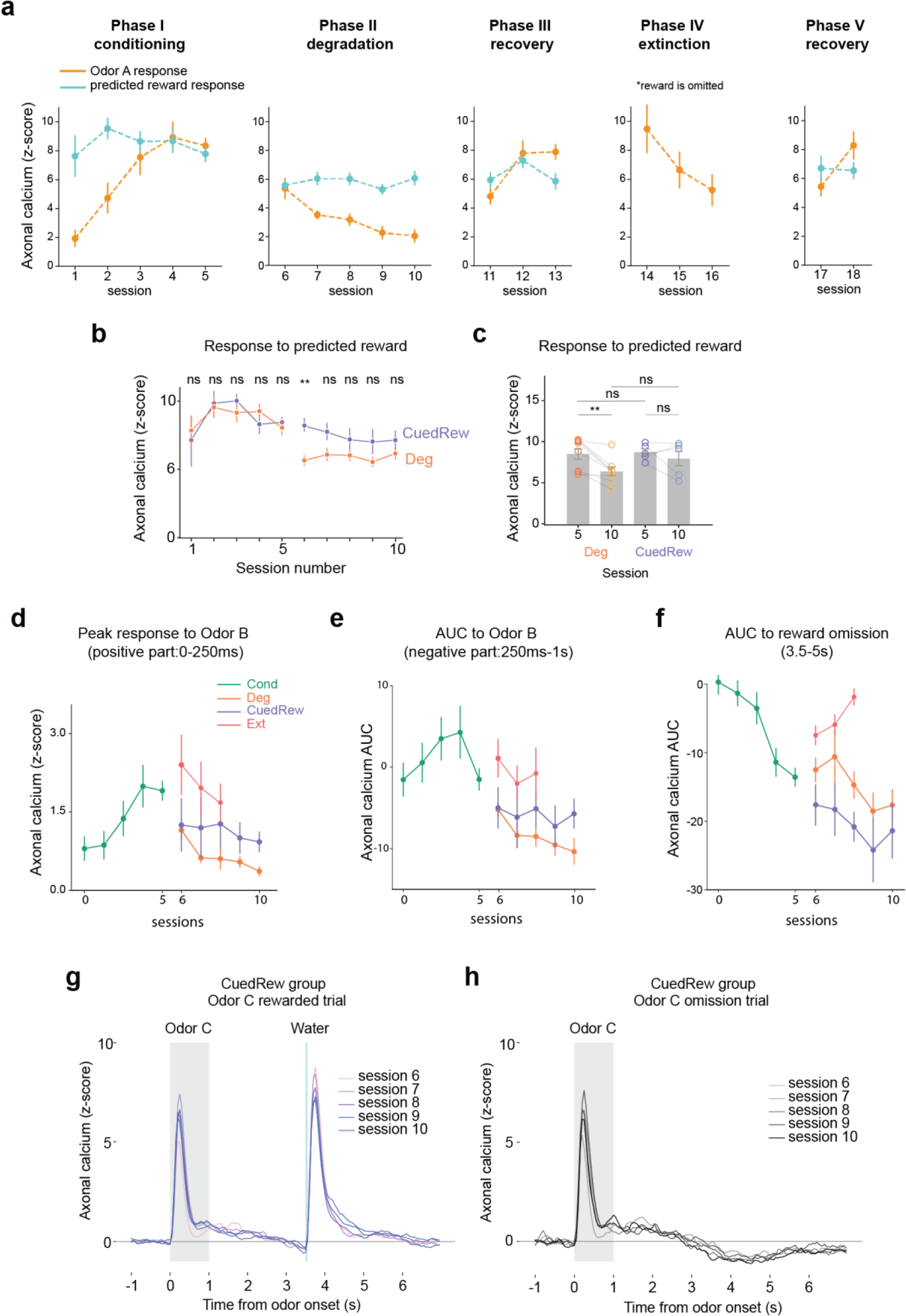
| Population Average Dopamine Response per session. (a) Mean peak dopamine axonal signal (z-scored) of cue response (orange) and reward response (cyan) in Odor A rewarded trial by sessions for the Deg group across multiple phases: Conditioning (Phase I), Degradation (Phase II), Recovery (Phase III), Extinction (Phase IV), and post-Extinction Recovery (Phase V). Error bars are SEM. (b) Mean peak dopamine axonal signal (z-scored) of reward response in Odor A trials by sessions for the Deg group (orange) and the CuedRew group (purple). Error bars are SEM. ns *P* < 0.05, \*\**P* < 0.001, Student’s *t*-test. (c) Mean peak dopamine axonal signal (z-scored) for the last session in Phase 1 and 2 for both Deg and CuedRew groups. Error bars represent SEM. ns, *P* >0.05; ***, *P* < 0.001, paired *t*-test. (d, e, f) Mean peak dopamine axonal signal (z-scored) across sessions for four distinct conditions, represented for various events. (g) Response to Odor C (rewarded) and (h) Odor C (omission), population average per session

**Extended Data Fig. 4.**
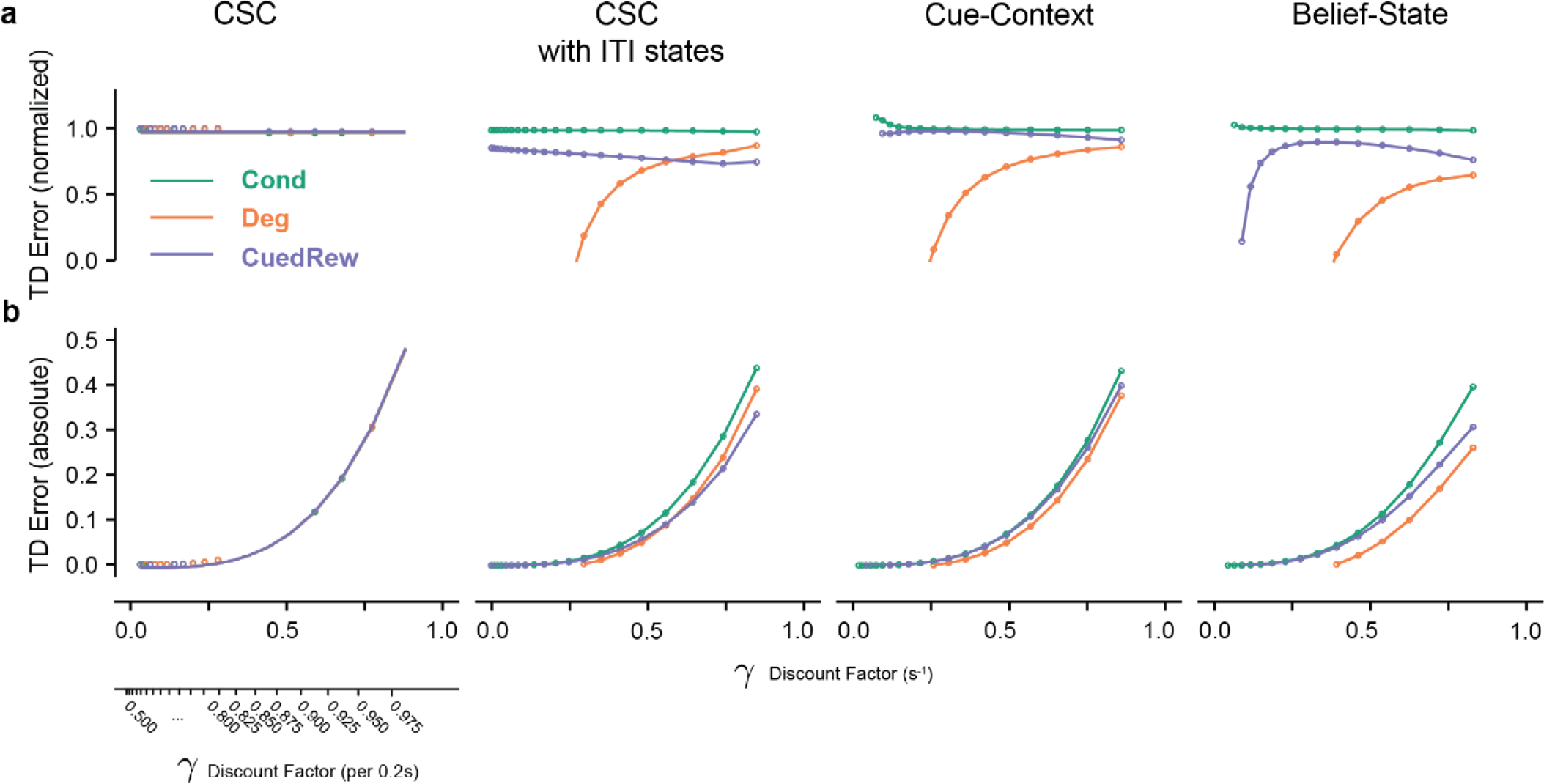
| Discount Factor determines modeled contingency degradation effect size. Influence of discount factor (γ) on relative predicted odor A response relative to Conditioning (a) or absolute (b), where reward size = 1 for four models presented in Figure 3. Bottom right scale showing discount factor converted to step size (0.2s), other axes use per second discount. Tested range: 0.5-0.975 discount per 0.2s in 0.025 steps.

**Extended Data Fig. 5.**
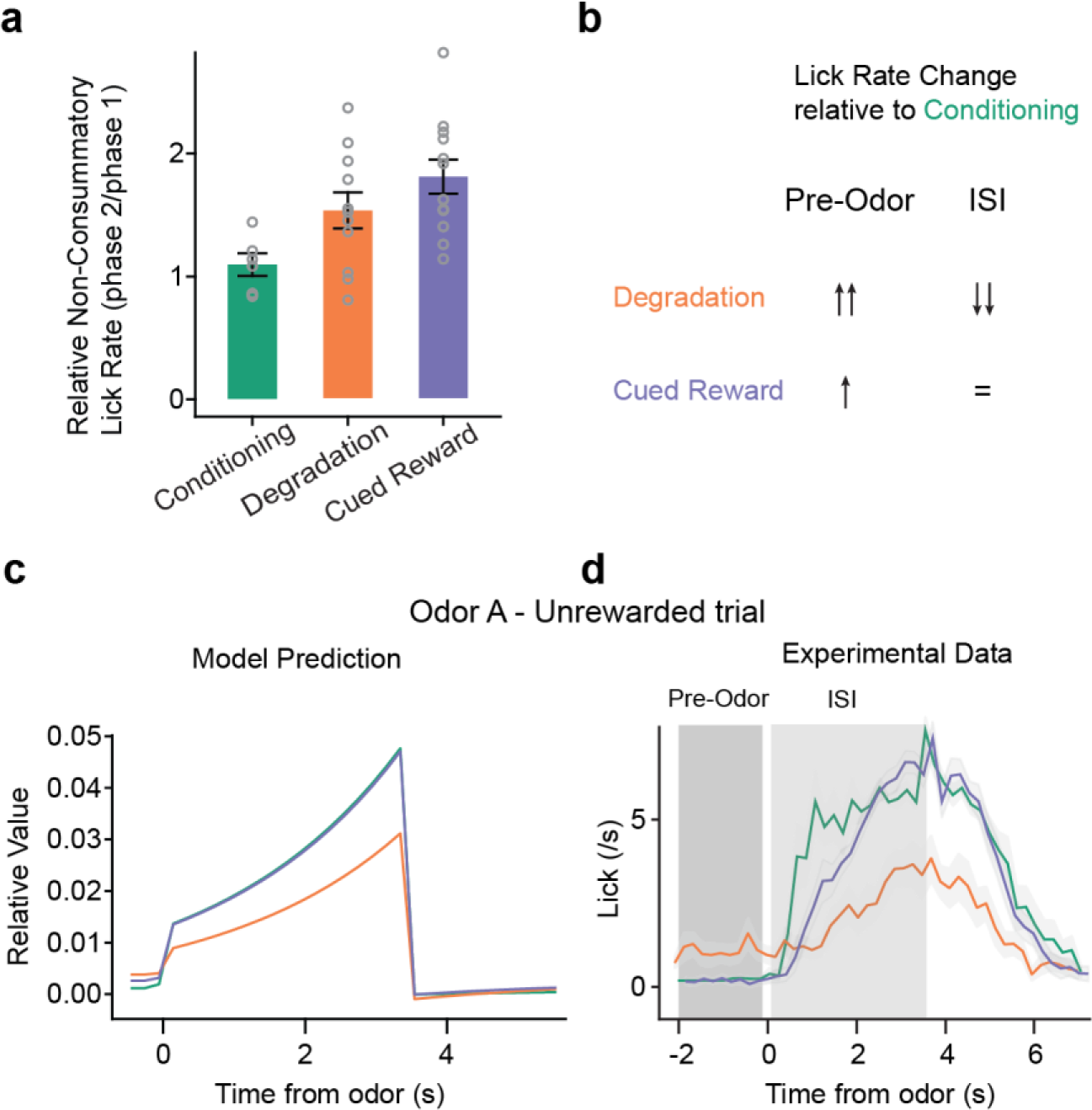
| Relative value explains decreased anticipatory licking during ISI during contingency degradation. (a) If each lick carries a small, fixed effort cost, a rational agent will lick proportionally to the total amount of rewards^75,76^. Plot show mean non-consummatory lick rate normalized to the Conditioning phase, suggesting that the Degradation and Cued Reward conditions elicit approximately twice the lick rate of the Conditioning condition, and thus proportional to the total reward quantity. Consummatory licks were considered any licks occuring in the 2 seconds following reward delivery. (b) Summary of lick rate changes relative to the Conditioning phase during the pre-odor period and the inter-stimulus interval (ISI). (c) Average relative value (current value/session total value, scaled by total reward) during odor A trial derived from the Belief-State model. Relative value, which is increased in the pre-odor period and thus decreased during the ISI, accounts for the change in licking pattern during unrewarded (and thus without consummatory licks) odor A trials. (d) Experimental data showing the actual lick rates recorded during Odor A unrewarded trials, compared over time, which aligns with the assumptions and predictions made in a,b, and c.

**Extended Data Fig. 6.**
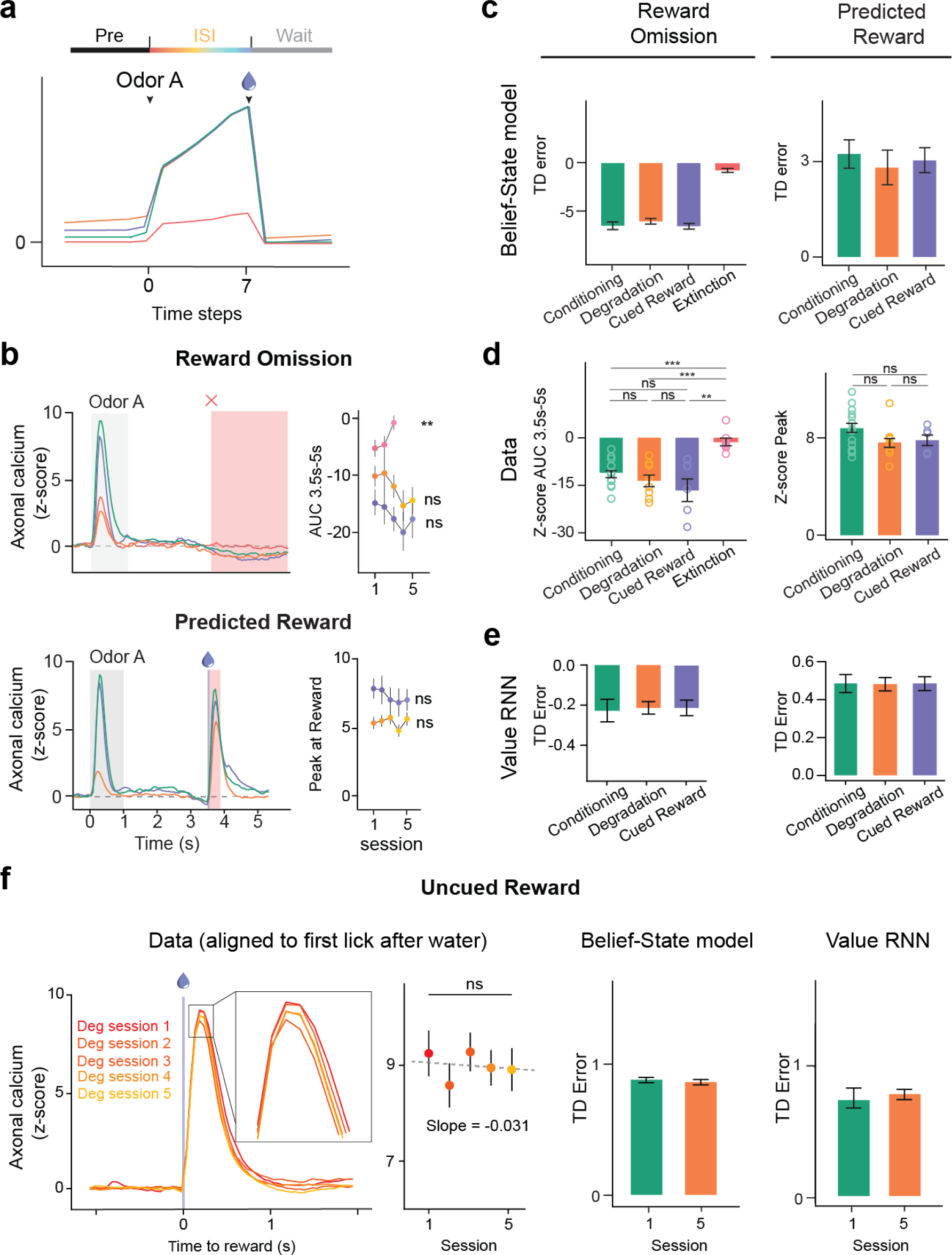
| Comparison of reward and omission responses between experimental data, Belief-State model and value-RNN predictions. (a) Plots averaged from one representative simulation of Odor A rewarded trial (*n* = 4,000 simulated trials) for four distinct conditions using the Belief-State model. Graphs are for the corresponding value function of Odor A rewarded trials, with Pre state, ISI state and Wait state annotated. (b) Z-scored DA axonal signals to reward omission and predicted reward following Odor A quantified from the red shaded area. Line graphs (right) shows mean z-scored response over multiple sessions for each condition. Statistical analysis was performed on data from the first and last session of these conditions. Error bars are SEM. ns, *P* > 0.05; **, *P* < 0.01, paired *t*-test. (c) The predictions of the Belief-State model for reward omission and predicted reward (mean, error bars: SD). (d) The experimental data for reward omission and predicted reward (mean, error bars: SEM). ns, *P* > 0.05; **, *P* < 0.01; ***, *P* < 0.001, Welch’s *t*-test. (e) The predictions of the Value-RNN models for reward omission and predicted reward (mean, error bars: SD). (f) The experimental data, TD error prediction by Belief-State model and Value-RNN model for uncued reward response in Degradation condition. While the Belief-State model captured the downward trend in response magnitude, none of the three statistical tests showed significant changes.

**Extended Data Fig. 7.**
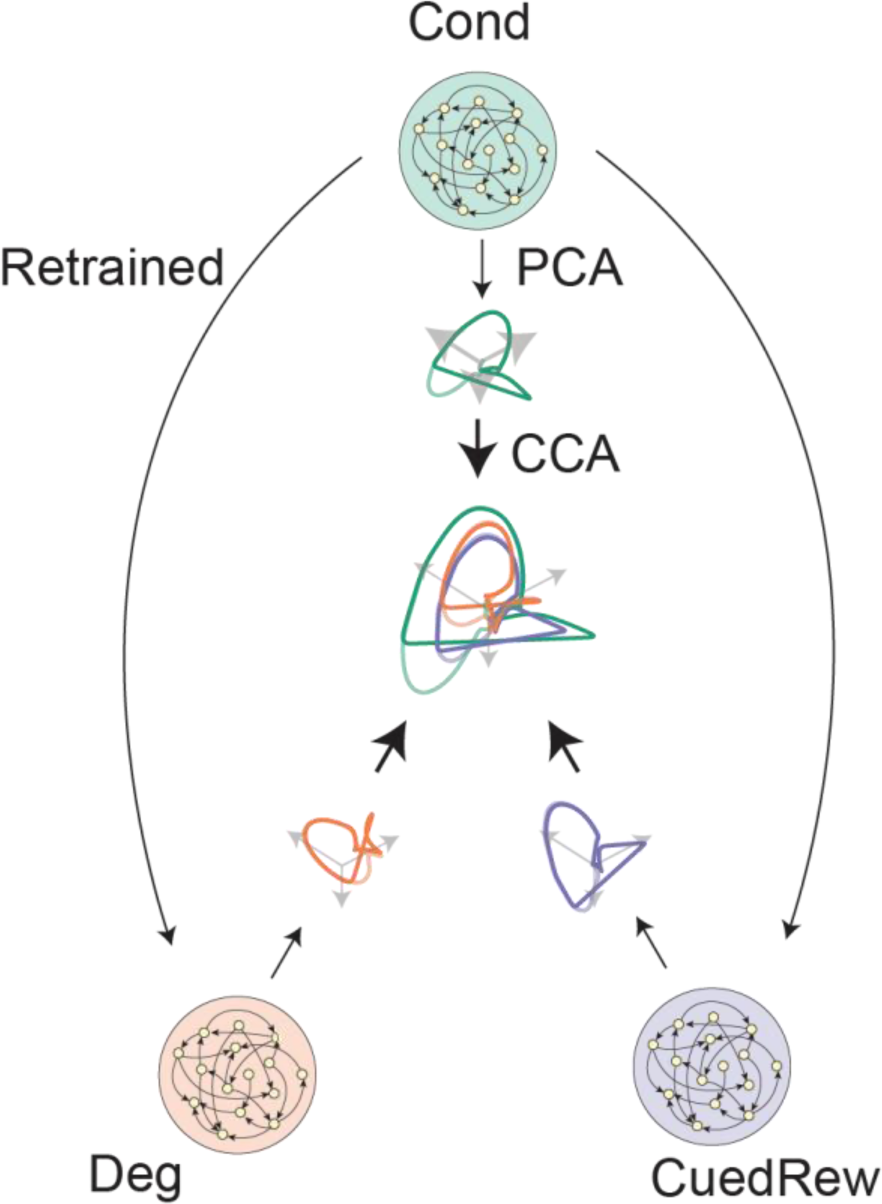
| Methodology for visualizing state space from hidden unit activity. Illustration for visualizing common state space of RNN models. RNN hidden unit activity was first projected into principal component space, then canonical correlation analysis was used to align between different conditions.

**Extended Data Fig. 8.**
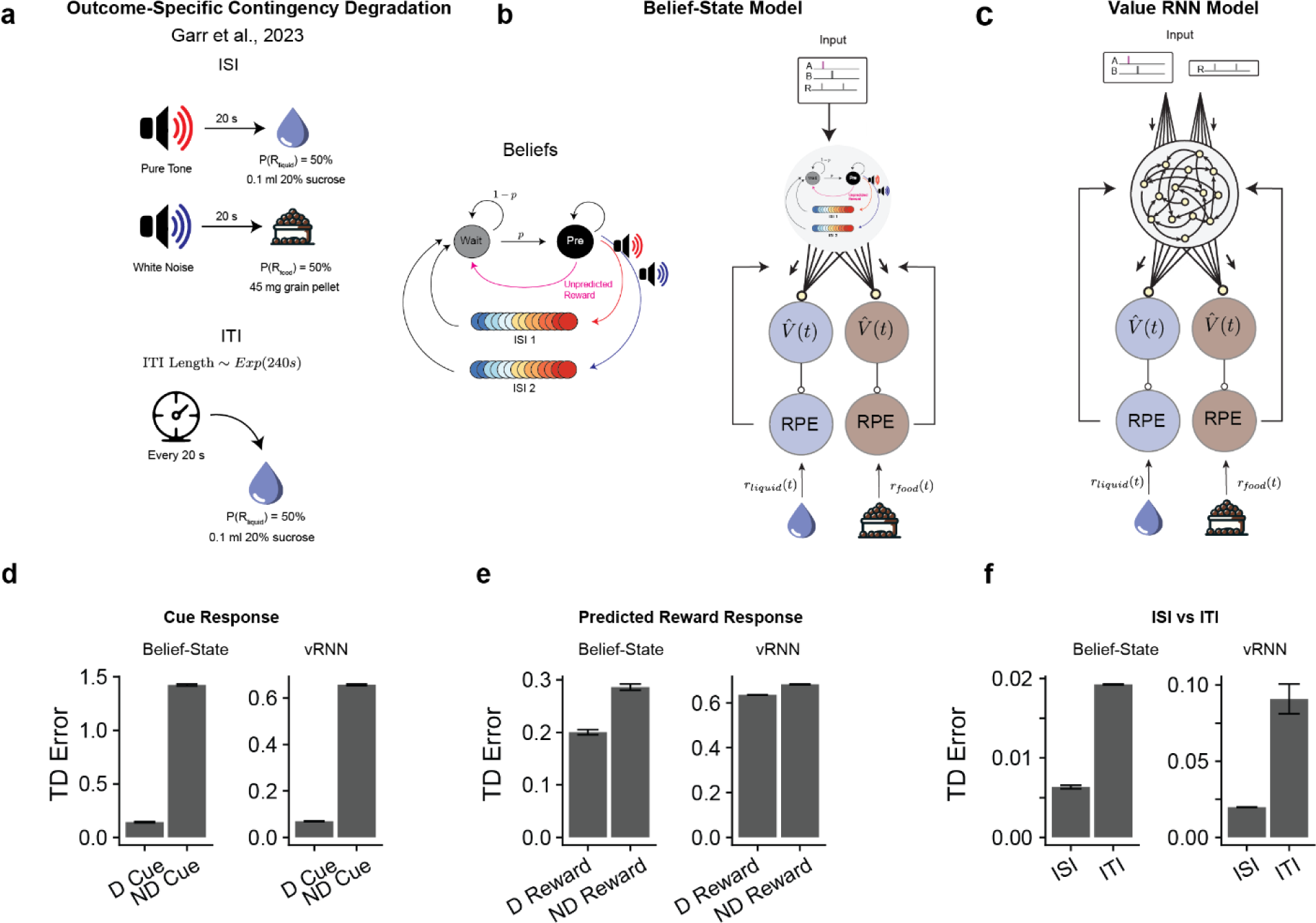
| Outcome-specific contingency degradation explained by Belief-State model and Value-RNN model. (a) Experimental design of Garr et al., two cues predicted either a liquid or food reward. During degradation, every 20 s the liquid reward was delivered with 50% probability. The ITI length was drawn from an exponential distribution with mean of 4 minutes. (b) Belief-State model design. The Belief-State model was extended to include a second series of ISI substates to reflect the two types of rewarded trials. The model was then independently trained on the liquid reward and food reward. (c) The value-RNN model design – as (b) but replacing the Belief-State model with the value-RNN, using a vector-valued RPE as feedback, with each channel reflecting one of the reward types. (d-f) Summary of predicted RPE responses from Belief-State Model and Value-RNN (vRNN). The RPE was calculated as the absolute difference between the liquid RPE and food RPE. Other readout functions (e.g. weighted sum) produce similar results. Both model predictions match experimental results with degraded (D) cue (panel d) and degraded reward (e) having a reduced dopamine response versus non-degraded (ND). Furthermore, average RPE during ISI (3 seconds after cue on) and ITI (3 seconds before ITI) capture measured experimental trend. Error bars are SEM.

## Extended Data Figures

**Extended Data Video 1: State Space trajectories**

Animation of trajectories in CCA space from RNN presented Figure 6e. In sequence, trajectories showing Odor A (rewarded), Odor A (unrewarded), Odor A (Rewarded and Unrewarded), Odor B and then all at once for the three conditions. Real time speed multiple indicated top right. ITI length is extended from training/actual experiment to demonstrate the return to original (‘Pre’) state in Conditioning and Cued Reward but the delayed return in Degradation condition.

